# Primary cilia loss promotes reactivation of morphogenesis and cyst-fission through a deregulated TGF*β*-ECM-Integrin axis in polycystic liver disease

**DOI:** 10.1101/2022.04.08.487546

**Authors:** Scott H Waddell, Yuelin Yao, Paula Olaizola, Edward J Jarman, Kostas Gournopanos, Ersi Christodoulou, Philippe Gautier, Joost PH Drenth, Timothy J Kendall, Jesus M Banales, Ava Khamseh, Pleasantine Mill, Luke Boulter

**Author notes:** These authors contributed equally to this work (Scott H Waddell, Yuelin Yao).

## Abstract

Pathological liver cysts are an important comorbidity in multiple diseases and syndromes(1, 2) driven by dysfunction of the primary cilium (PC), a complex sensory organelle that protrudes from the apical surface of biliary epithelial cells (BECs)(3, 4). The essential nature of PC in liver development(5, 6) makes understanding the molecular role of this organelle in the structural maintenance of the adult bile duct challenging. Here, we show that PC loss deletion of *Wdr35* in adult mouse BECs is sufficient to cause bile duct expansion, driving cyst formation through the *de novo* production of a fibronectin-rich pro-cystic microenvironment. This newly formed niche promotes both cell-autonomous changes in cell shape and duct-level mechanical rearrangements that converge to drive cyst-fission, a novel process whereby single, large cysts undergo morphological splitting. This process gives rise to many, smaller polycystic progeny and can be halted by pharmacological inhibition of a specific pro-cystic integrin receptor.

## Results

Genetic alterations in PC genes result in hepatorenal fibrocystic diseases (HRFCD), such as hereditary polycystic kidney diseases, which is caused by mutations in *PKD1*, *PKD2* or PKDH1(7–9). Interestingly, isolated polycystic liver disease (PCLD) is driven by mutations *SEC63*, *GANAB* or *PRKCSH*, among others, all of which encode ER-associated proteins that play roles in protein transport, folding and trafficking to PC(10, 11). Complex fibropolycystic features are also associated with syndromic ciliopathic disease in which key PC genes, such as intraflagellar transport (IFT) genes (*WDR35/IFT121*, *WDR19/IFT144* and *IFT56*) that are required to build and maintain PC are mutated(12–14). Across these diseases, cysts form throughout the liver, however it remains unclear whether BEC PC-loss *per se* in the adult is sufficient to promote liver cystogenesis (**Figure 1a** summarises the mutations associated with PCLD).

**Fig. 1.**
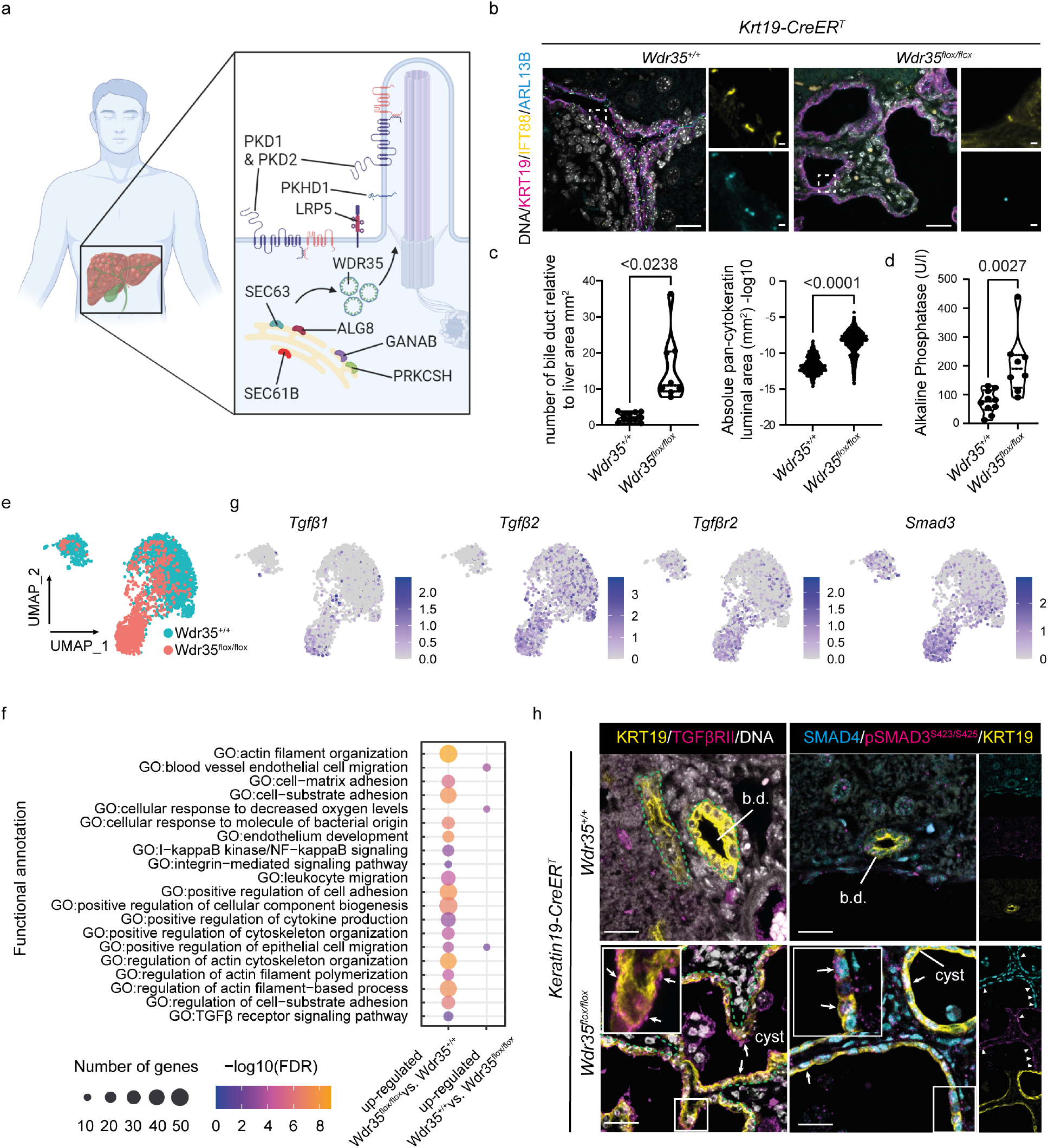
Selective loss of *Wdr35* in BECs results in PC loss and the formation of polycystic liver disease linked to TGF*β* signalling. **a.** The genetic mutations associated with autosomal polycystic kidney disease (in which cysts arise in kidney and liver), isolated polycystic liver disease and syndromic cystic diseases (CHF) can affect the formation, stability or protein trafficking to the primary cilium. **b.** Cilia are defined by the presence of ARL13B (cyan) and AcTUB (yellow) expressed by KRT19 expressing bile ducts (magenta), DNA in grey. Scale bar = 100 *μ*m. **c.** Number (*Wdr35*^+/+^ N=11 animals and *Wdr35*^−/−^ N=8 animals) and area of ducts or cysts in 12 month *Wdr35*^+/+^ and *Wdr35*^−/−^ livers (*Wdr35*^+/+^ n=812 ducts and Wdr35−/− n=3834 cysts). **d.** Serum alkaline phosphatase (ALP) levels of 12 month *Wdr35*^+/+^ (N=10 animals) and *Wdr35^−/−^*) (N=8 animals) mice. **e.** UMAP of single cells showing segregation of *Wdr35*^+/+^ (cyan) and *Wdr35*^−/−^ (salmon) cells. **f.** Go term analysis of marker genes in cluster 1 (87% composed of *Wdr35*^+/+^ cells) and cluster 2 (97% composed of *Wdr35*^−/−^ cells), identified by hiercharcical clustering. **g.** UMAPs of normal (*Wdr35*^+/+^ n=3060 cells) and cystic cells (n=966 cells) showing single cell expression of *Tgfβ1*, *Tgfβ1*, *Tgfβr2* and *Smad3*. **h.** Immunohistochemistry of control tissue and *Wdr35^−/−^* cyst bearing livers stained for KRT19 (yellow), TGFBRII or pSMAD3^*S*423*/S*425^ (magenta, white arrows denote positive staining in BECs) and SMAD4 (cyan). DNA is represented in grey. Green dotted lines denotes the boundary of the duct or cyst, white arrows denote positive cells (scale bar=100 *μ*m).

Previous reports suggest that in liver cysts from autosomal dominant polycystic kidney disease, PC length decreases with cyst size(15). To determine whether loss of a functional primary cilium was sufficient to promote the formation of cysts within the liver we deleted the *Wdr35/Ift121* gene specifically in BECs (using *Krt19-CreER^T^* mice, **Supplementary figure 1a-1b**). WDR35/IFT121 is part of a highly conserved, non-core IFT-A complex that we have shown is essential for the transport of ciliary membrane cargo, and necessary for PC elongation and signalling(16). This approach enables us to define the roles of PC specifically in adult biliary homeostasis, separating whether changes in PC *per se* cause PCLD rather than being a consequence of errant ductal plate patterning found in embryonic ciliamutants(6, 17). In control *Krt19-CreER^T^;Wdr35*^+/+^ (referred to as *Wdr35^+/+^*) bile ducts, IFT88/ARL13B-positive PC protrude from the apical surface of BECs; however in *Krt19-CreER^T^;Wdr35*^−/−^ (abbreviated to *Wdr35*^−/−^) BEC PC are largely absent and instead exist as IFT88-positive, shortened rudiments lacking ARL13B and normal PC function, as we have previously described (**Figure 1b** and **Supplementary figure 1c**).

In *Wdr35^+/+^* mice, bile ducts are arranged around portal veins forming a network of small, pan-cytokeratin positive ductules. Following deletion of *Wdr35* from BECs, pancytokeratin positive ducts become progressively polycystic between 6 and 12 months, with a small but significant increase in BEC proliferation. This results in an increase in both the number of cystic ducts and their area over this period (**Figure 1c** and **Supplementary figure 1d-1f**) similar to reports for adult *Pkd2* deletion (17). The development of PCLD following *Wdr35-loss* also results in an elevation of serum alkaline phosphatase levels, indicating that PC-loss and the formation of liver cysts leads to biliary inflammation (**Figure 1d**) without affecting other hepatic functions (**Supplementary figure 2**).

Whilst symptomatic liver cysts are a comorbid factor in multiple diseases, little is known about the molecular processes that enable normal ducts to become pathological cysts. We therefore sought to define the molecular changes that BECs undergo as they become cystic. CD45-/CD31-/EpCAM+ BECs from *Wdr35*^+/+^ or *Wdr35*^−/−^ livers were isolated 12 months following PC-loss (**Supplementary figure 3**). Using 10X single cell RNA sequencing (scRNA) we analysed the gene expression of 3060 normal and 966 cystic cells. BECs cluster into 4 distinct and highly stable populations (**Figure 1e** and **Supplementary figure 4a-d**) based on hierarchical clustering and whilst normal and cystic cholangiocytes cluster separately at the transcriptional level. RNA velocity (18, 19) predicts that cystic cells are derived from normal BECs as expected rather than transformation from other hepatic cell types (**Supplementary figure 4e**). Within the four clusters identified, the majority of *Wdr35*^+/+^ cells fall within cluster 1, however most *Wdr35*^−/−^ cells occupy cluster 2 (cluster 3 and 4 are a mixture of cells from both WT and mutant animals) and regardless of genotype cells continue to express a number of BEC-specific markers (**Supplementary figure 5a** and **5b**). A large number of genes were differentially expressed between clusters 1 and 2 (**Supplementary table 1** and **Supplementary figure 6a** and **6b**) and gene ontology (GO) term analysis identified multiple transcriptional signatures that were enriched in *Wdr35^−/−^* cells, including Ca2+ and MAPK signalling, which have been associated with the formation of renal(8) and hepatic(20) 2022 cysts (**Supplementary figure 7a-7d**). Hedgehog signalling, which is tightly controlled by the cilium in other contexts (21, 22), was not altered in cystic cells (**Supplementary figure 7e**); rather, GO terms associated with cytoskeleton remodelling and cell-matrix adhesion (23, 24), namely TGF*β* (25)(and integrin(26) signalling were changed (**Figure 1f** and **Supplementary table 2**) adding further support to the idea that alterations in the extracellular matrix (27) and cytoskeletal remodelling, in addition to induction of known cystogenic signalling signatures promote the generation of cysts *in vivo.*

Canonical TGF*β* promotes SMAD3 phosphorylation, and with SMAD4 regulates inflammatory cytokines and extracellular matrix gene transcription. In the liver, TGF*β*-SMAD signalling is a central driver of biliary fibrosis (28), but is constrained to nascent portal fibroblasts surrounding ducts that produce collagen in response to TGF*β*(29). In contrast, our data suggests that cystic BECs themselves may be targets of TGF*β* ligands. We sought therefore to define the role of TGF*β* in liver cyst formation. In addition to upregulation of *Tgfb1* and *Tgfb2* ligands, cystic BECs significantly upregulate transcription of the receptor *Tgfbr2* and the downstream effector of this pathway *Smad3* (**Figure 1g**). We confirmed levels and localisation by immunofluorescence whereby *Wdr35^+/+^* BECs do not express TGFBRII; however 12-months following Wdr35-loss, TGFBRII is apically enriched on cystic BECs (**Figure 1h**). To test whether the presence of TGFBRII on cystic BECs was indicative of cysts being TGF*β* responsive, we examined SMAD expression. In control *Wdr35*^+/+^ bile ducts, SMAD4 is present but lowly expressed and with the exception of small numbers of cells, (active) pSMAD3^*S*423/*S*425^ is absent. In contrast, in cystic BECs, high nuclear SMAD4 and extensive pSMAD3^*S*423/*S*425^ staining is observed. Active SMAD signalling in the epithelium of ADPKD renal cysts has been described (30) suggesting there could be common TGF*β*-mechanisms of cyst formation between diseases. To test this, we assessed cystic tissue from PCLD patients where either the causative mutation has not been identified or which are driven by mutations in *SEC63* (an ER protein thought to be involved in processing of PC proteins amongst others(31)) or *PRKCSH* (an essential subunit of an ER-residing glucosidase, necessary for PKD1 trafficking(32)) (**Figure 1a**). In these genetically diverse human pathologies, elevated nuclear SMAD4 and pSMAD3^*S*423*/S*425^ are detected in cystic BECs, confirming that TGF*β* signalling is a common hallmark of liver cyst formation (**Supplementary Figure 8a**).

As TGF*β* signalling is activated in cystic BECs, we postulated that TGF*β*-SMAD is necessary for the conversion of normal ducts into cysts. We adapted previous liver organoid methodologies(33) to isolate bile ducts from wild type mice. These *ex vivo* ducts are cultured in Matrigel and form rounded sphere-like cystic structures within 72 h (**Figure 2a**). In this assay, treatment of ducts with the SMAD3-inhibitor SIS3 significantly reduces the area of cysts, and rather than developing into dilated cystic structures, ducts remain as tubes with closed ends (**Figure 2b**). Given the failure of SIS3-treated ducts to form cysts and evidence from our BEC-specific scRNA data (**Figure 1**) we hypothesised that signalling through SMAD3 enables BECs to alter their physical microenvironment, thereby promoting cystogenesis. To define this, we isolated total protein from freshly isolated bile ducts or ducts cultured for 72 h and treated with either SIS3 or a vehicle alone. Using a custom Reverse Phase Protein Array, a sensitive antibody-based proteomic approach biased to detect changes in ECM and cell-adhesion proteins, we found that when compared to freshly isolated ducts, 72 h cultured ducts increase fibronectin production as they become cystic. Furthermore, the level of fibronectin produced is significantly reduced in cultures that have been treated with SIS3 (**Figure 2c**).

**Fig. 2.**
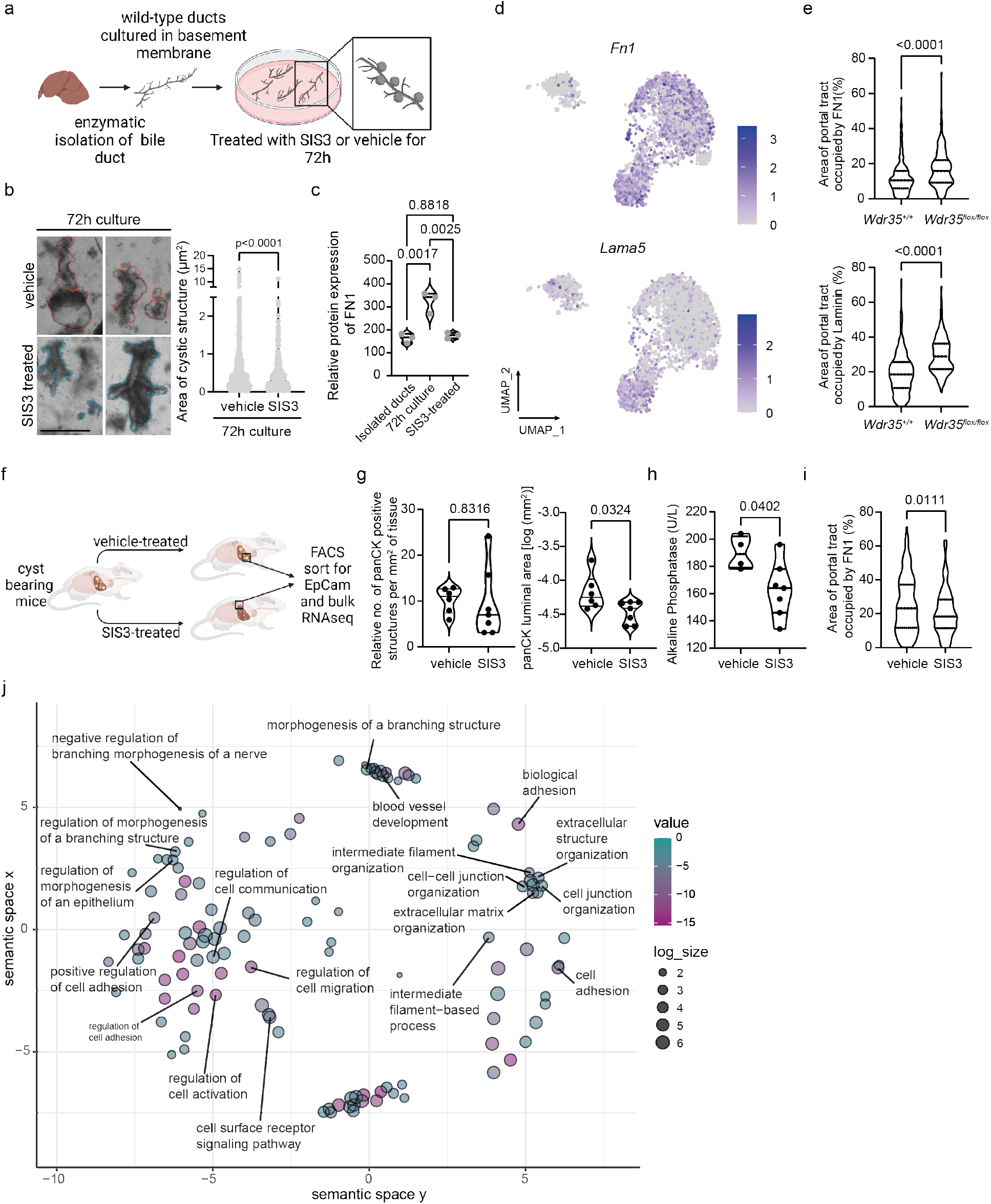
Cystic epithelial cells are sensitised to TGF*β* signalling thereby promoting the formation of a pro-cystic ECM. **a.** Schematic representation of duct-to-cyst cultures where wild-type bile ducts are plated in ECM and form cysts within 72 h of culture and can be treated with small molecule modulators like SIS3. **b.** Images, left, show typical structures of ducts and organoids in 72 h culture with and without SIS3 treatment. Quantification, right, of *ex vivo* cultures treated with the SMAD3-inhibitor, SIS3 (n=487 ducts in vehicle and n=378 ducts in SIS3). **c.** Protein expression of fibronectin in freshly isolated ducts, after 72h culture with vehicle treated duct-to-cyst cultures or SIS3-treated duct-to-cyst cultures (n=3 where each n represents ducts isolated from two separate mice per condition). **d.** UMAP showing single cell transcript expression of *Fn1* and *Lamb3*. **e.** Quantification of immunohistological expression of fibronectin (n=284 *Wdr35*^+/+^ cysts from 11 animals and n=237 *Wdr35*^−/−^ from 8 animals) and pan-laminin (n=177 v cysts from 11 animals and n=299 *Wdr35*^+/+^ cysts) around bile ducts and cysts in 12 month livers. **f.** Schematic detailing the protocol for treatment of cyst bearing *Wdr35*^−/−^ mice with SIS3. **g.** Number and log size of panCK positive structures in cyst bearing mice treated with vehicle alone or SIS3 for 3 weeks (N=6 animals in vehicle group and N=7 animals in SIS3 treated). **h.** Serum Alkaline Phosphatase levels in cyst bearing mice treated with vehicle (N=4 animals) or SIS3 (N=7 animals). **i.** Fibronectin levels in *Wdr35*^−/−^ animals treated with SIS3 (n=277 cysts from 6 mice) or vehicle (n=196 cysts from 7 mice) alone. **j.** REViGO output showing the rationalised GOterms following GOrilla analysis.

The fibronectin gene, *Fn1* is a target of SMAD3 *in vitro*(34) and our data from both mouse (**Figure 1**) and human cells (**Supplementary figure 8c**and **8d**) indicates that via TGF*β*-SMAD3, BECs alter their own microenvironment by inducing fibronectin expression, in addition to a range of other ECM components. Fibronectin acts as a nucleating protein for fibrillar collagens (including collagen-I and collagen-IV), which interact with laminins to promote formation of the basement membrane(35). *Wdr35^−/−^* cystic BECs express high levels of *Fn1*, *Lama5*, *Lamb3*, *Lamc1*, *Col4a1* and *Col4a2*, but not *Col1a1* or *Col1a2* when compared to *Wdr35*^+/+^ BECs (**Figure 2d**; **Supplementary figure 9a** and **9b** and **Supplementary table 1**). In addition to these transcriptional changes, we confirmed the ECM surrounding *Wdr35^−/−^* cysts is different from *Wdr35^+/+^* bile ducts, showing an expanded basement membrane with increased fibronectin and laminins (**Figure 2e**).

Our *ex vivo* duct culture shows BEC-derived fibronectin promotes cyst expansion via SMAD3, therefore we administed 6-month cyst-bearing *Wdr35^−/−^* animals the SMAD3 inhibitor, SIS3 or vehicle alone for 3 weeks (**Figure 2f**). Inhibition of SMAD3 did not reduce the number of panCK-positive cysts in the *Wdr35^−/−^* liver; however, it significantly reduced cyst size (**Figure 2g**). Moreover, SMAD3-inhibition ameliorates biliary alterations as evidenced by reduction of levels of serum alkaline phosphatase (**Figure 2h**) and glutamate dehydrogenase (**Supplementary figure 10a**) and is associated with a significant reduction in fibronectin deposition around cysts following SIS3-treatment (**Figure 2i**). Finally, we took an agnostic approach to examine how TGFβ-SMAD3 regulates cyst formation in PCLD. CD45-/CD31-/EpCAM+ BECs were isolated from *Wdr35^−/−^* cysts treated with either SIS3 or vehicle and subjected to bulk RNA sequencing. Upon SIS3-treamtent, cystic BECs were transcriptionally distinct from those isolated from vehicle animals (**Supplementary table 3**). Analysis with GOrilla and REViGO (**Supplementary table 4**) showed that SIS3-treatment changes gene sets associated with tubular morphogenesis, branching, and organisation of the ECM, reiterating that SMAD3 regulates a range of biological processes required for cyst generation (**Figure 2j**).

How cystic BECs receive and interpret signals from this changing extracellular niche remains unclear. In heterocellular scRNA data, receptor-ligand interaction modelling predicts functional interactions between different cell-types(36, 37). Here, using SingleCellSignalR(36) we sought to define whether there are cell-type autonomous signals between BECs and address whether these interactions change as BECs transition to cysts. Within wild-type BECs, a number of receptor-ligand interactions were identified (**Figure 3a** and **Supplementary table 5**), notably between Fn1, encoding for fibronectin, and *Itgb1 (β* 1-integrin), a canonical ECM receptor which heterodimerises with other *α*-integrin subunits to sense alterations in the ECM. Analysis of cystic *Wdr35^−/−^* BECs also identified candidate interactions between *Itgb1* and Fn1, however this *Fn1* domain was expanded including putative *Wdr35^−/−^*-specific interactions with *Itga2*, *Itga3*, *Itgav*, *Itgb6*, *Cd44* and *Plaur* (**Figure 3a** and **Supplementary Table 6**). As integrin-receptor affinity is altered depending on the relative expression of both *α-* and *β*-subunits(38), we clustered wild-type and cystic BECs based on genotype (*Wdr35^+/+^* vs *Wdr35^−/−^*) to define which integrin subunits are most differentially expressed between these groups (**Supplementary Figure 11a**). *β*1-integrin (*Itgb1*) is universally expressed by BECs whether they are normal or cystic, whereas *α*2-integrin (*Itga2*) is selectively expressed by *Wdr35^−/−^* mutant cells (**Figure 3b** and **Supplementary figure 11b**). Integrin-*α*2*β*1 is a relatively promiscuous integrin that binds laminins, fibrillar collagens and fibronectin and both ITGB1 and ITGA2 proteins are expressed on cystic BECs from *Wdr35^−/−^* mice (**Figure 3c**, **Supplementary figure 11b**). Furthermore, downstream mediators of integrin signalling Pxn, *Fermt1, Ilk, Tln1, Actn1* and *Rhoc* were also upregulated in *Wdr35^−/−^* cysticBECs (**Figure 3d**). In other ductular systems, ECM-integrin signalling regulates the actin cytoskeleton thereby modulating cellular width and apico-basal tension((39–41). In liver cysts, BECs have a generalised increase phosphorylated myosin light chain 2 (pMLC2) expression compared to their wildtype counterparts (**Figure 3e-3g**). In *Wdr35^+/+^* BECs, low levels of pMLC2 are preferentially localised to the basal domain of the cell, however cystic BECs have significantly increased apical levels of pMLC2 (**Figure 3h**). This increase in apical pMLC2 levels in *Wdr35^−/−^* cystic BECs is associated with a significant increase in BEC width (from lateral membrane to lateral membrane, **Figure 3i**). Together, our data support that loss of PC signalling in BECs lead to profound cell-autonomous morphogenetic changes, driving cyst formation in the adult liver (**Figure 3j**).

**Fig. 3.**
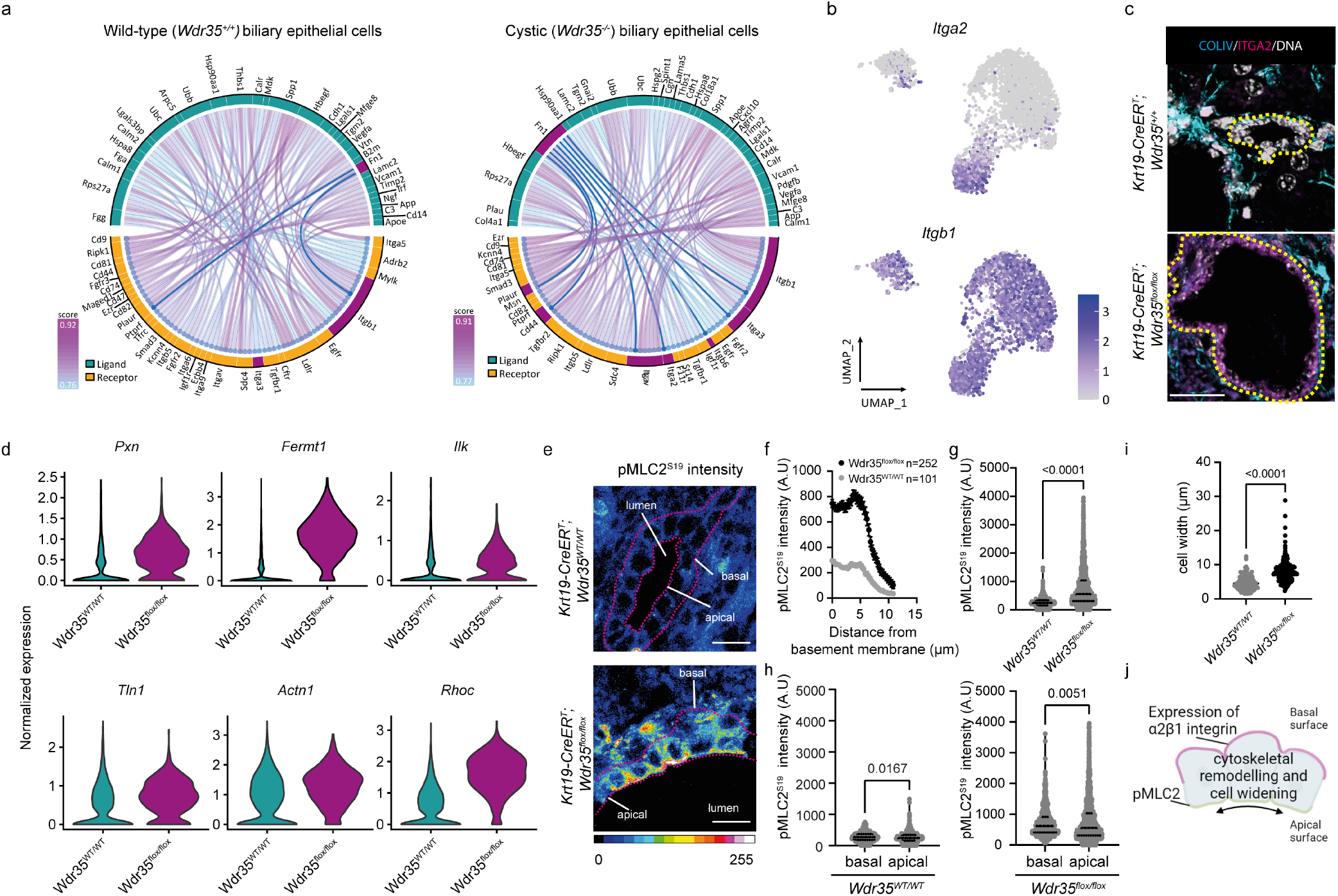
Cystic cells expand their ability to interact with a fibronectin-rich microenvironment and alter their cell shape. **a.** SingleCellSignalR analysis of ligand-receptor interactions in scRNAseq data from *Wdr35*^+/+^ and *Wdr35*^−/−^ animals. **b.** UMAP showing the transcriptional expression of *Itga2* and *Itgb1* **c.** Immunohistochemistry for Collagen-IV (cyan) and *α*2-integrin (magenta) in normal and cystic livers. Scale bar=100 *μ*m. Yellow dotted line denotes the boundary of a duct or cyst. **d.** Histograms from scRNA data of *Pxn*, *Fermt1*, *Ilk*, *Tln1*, *Actn1*, *Rhoc* comparing median transcript levels between *Wdr35*^+/+^ and *Wdr35^−/−^* biliary cells. **e.** Intensity projection of pMLC2 staining in normal bile ducts compared to cysts. Dotted line denotes boundary of ducts and cysts. **f.** Quantification of pMLC2 intensity across the apico-basal axis of normal biliary cells (grey line) and cystic epithelial cells (black line). **g.** Average pMLC2 intensity in normal and cystic biliary cells. **h.** Apical and basal pMLC2 levels in biliary cells from *Wdr35*^+/+^ and *Wdr35*^−/−^ animals (for **f-h**, n=101 *Wdr35*^+/+^ cells and n=252 *Wdr35*^−/−^ cells from 4 animals per group). **i.** Width of biliary cells from *Wdr35*^+/+^ (n=165 cells from 4 livers) and *Wdr35^−/−^* (n=172 cells from 4 livers) mice. **j.** Schematic demonstrating the relationship between integrin-high states and cytoskeletal rearrangements *Wdr35^−/−^* animals treated with SIS3 (n=277 cysts from 6 mice) or vehicle (n=196 cysts from 7 mice) alone.

As in our mouse model of PCLD, patients with cystic liver diseases have high levels ITGA2 and pMLC2 on cystic BECs regardless of underlying causative genetics (**Supplementary Figure 12a**), suggesting ciliary or trafficking-mediated disruptions converge on similar pathomechanisms. We further confirmed the presence of integrin-*α*2*β*1 in cystic epithelium from both liver and kidney cysts from the *Pkhd1*^pck^ rat (a model of autosomal recessive polycystic kidney disease, ARPLD; **Supplementary Figure 12b and 12c**), suggesting that a transition to integrin-*α*2*β*1 mediated-sensing of the microenvironment may be a common mechanism during cyst formation across genetic models, species and tissues.

Patients with PCLD, by definition, present with multiple (>10) cysts in their liver; however, it remains unclear whether each cyst originates clonally from a mutant precursor cells (i.e. many separate originating events) or whether cysts themselves divide to produce a polycystic pattern of disease (i.e. cells of origin are divided between multiple cysts). Recent work demonstrated that ductal plate malformation during embryogenesis and/or loss-of-heterozygosity is required for cyst formation(42–44). In other epithelial systems, loss-of-heterozygosity (in tumour suppressor genes for example) drives clonal selection(45, 46). To determine whether the cysts that form following *Wdr35*-loss are derived from multiple individual clonal events or because of the division of cysts, we crossed *K19CreER^T^;Wdr35^flox/flox^* mice with the Confetti lineage reporter mouse line (*K19CreER^T^*;*Wdr35^flox/flox^*;*R26^LSL-Confetti^*). In this model, upon CRE recombination, cells are stochastically labelled with one of nGFP, mCFP, cYFP or cRFP (or no fluorescence)(47), thereby enabling clonal tracing of recombined cells in BECs (**Figure 4a**). Six months following *Wdr35*-deletion (and Confetti labelling) livers were surveyed using 3D confocal imaging using the FUnGI protocol(48)). In *Wdr35*^+/+^-Confetti mice, confetti-labelled BECs remained as single cells (**Supplementary figure 13a**). In *Wdr35*^−/−^-Confetti mice, small, isolated cysts were clonal in colour i.e. derived from a single labelled BEC (**Figure 4b** and **Supplementary figure 13a**). However, where clusters of cysts had formed (i.e. regions of polycystic disease), they were comprised of labelled and unlabelled cells. Furthermore, within clusters of cysts, adjacent cyst walls often shared the same label colour, strongly implying that polycystic disease forms from multiple mutant cells and those large cysts may undergo cyst-fission and “pinch off” to generate smaller, daughter cysts (**Figure 4b**).

**Fig. 4.**
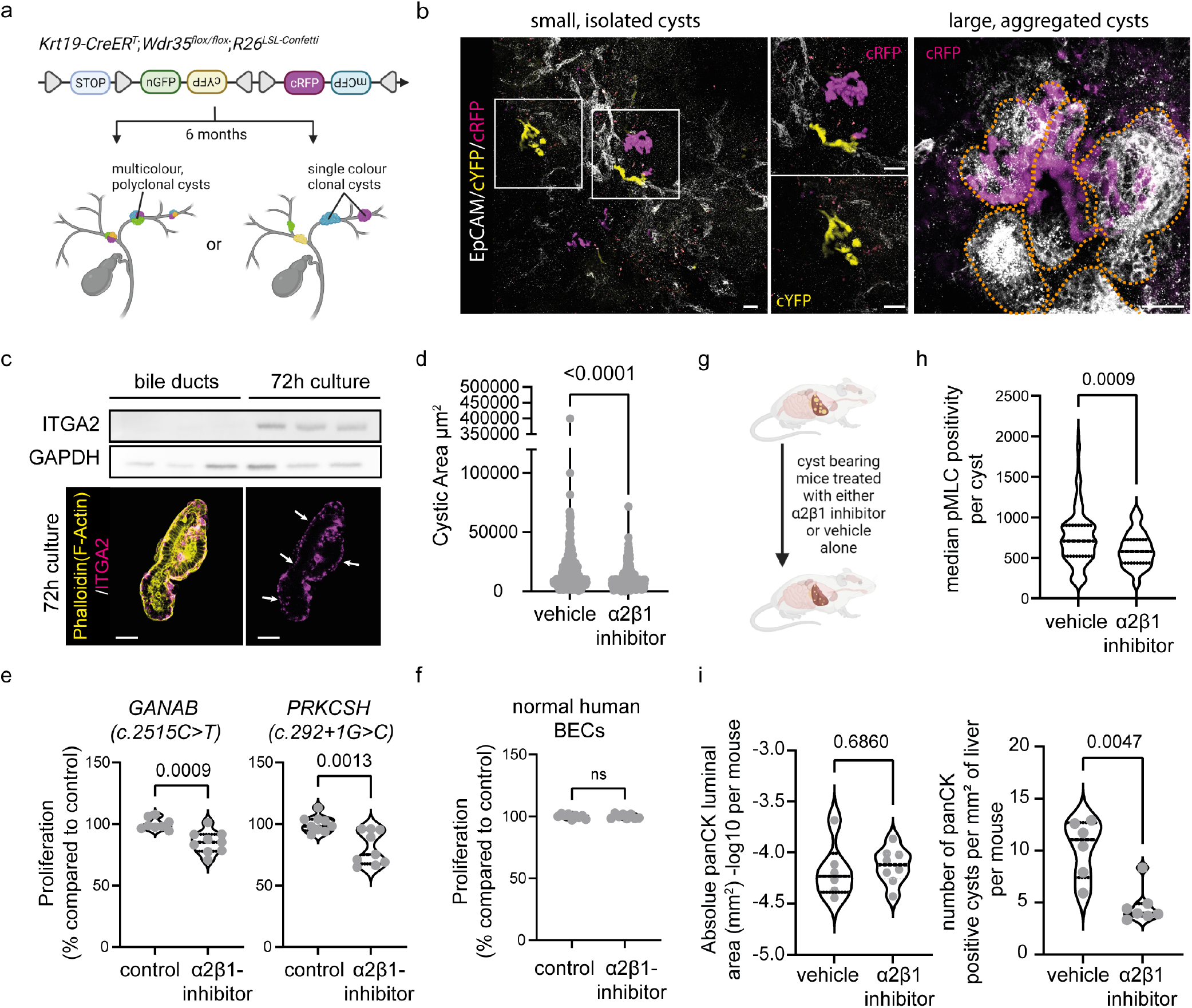
Integrin-*α*2,*β*1 promotes hepatic cystogenesis by mediating cyst fission. **a.** Schematic representation of the (*K19CreER^T^*;*Wdr35^flox/flox^*;*R26^LSL-Confetti^*) and the lineage traced outcomes of cysts in this model, where cysts could arise from either a single clone or from multiple clones. **b.** (*K19CreER^T^;Wdr35^−/−^;R26^LSL-Confetti^*) livers cleared with FUnGI and imaged for EpCAM (grey), cYFP (yellow) or cRFP (magenta). White boxes represent magnified regions of interest. Orange dotted line denotes a collection of cysts that share a single mutant clone. Scale bar = 100 *μm* **c.** Immunoblot showing ITGA2 (integrin-*α*2) and GAPDH protein expression in freshly isolated bile ducts compared to those cultured for 72 h. Immunocytochemistry shows the localisation of ITGA2 (magenta) and F-actin (phalloidin; yellow) in 72 h cultured ducts (white arrows highlights ITGA2 localized at the basal surface of the cell). **d.** Area of cultured ducts as they become cystic when treated with either the Íntegrin-*α*2*β*1 inhibitor TC-I 15 (n=439) or vehicle alone (n=354) for 72 h. **e.** Proliferation of BECs from patients with mutations in GANAB or PRKCSH treated with the Íntegrin-*α*2*β*1 inhibitor TC-I 15 or vehicle alone (N=9 represents experimental replicates). **f.** Healthy control human BECs treated the same way as (**e**). (N=9 represents experimental replicates). **g.** Schematic demonstrating the strategy for in vivo inhibition of Íntegrin-*α*2*β*1. **h.** Median expression of pMLC2 staining in cystic cells treated with either vehicle (n=176 cysts from 7 mice) or Íntegrin-*α*2*β*1 inhibitor, TC-I 15 (n=73 cysts from 6 mice). **i.** Quantification of cystic luminal area in the livers of mice treated with TC-I 15 (N=6 mice) or vehicle (N=7-8 mice) or number of cysts in the same experiment (right panel).

To explore this further, we adapted our *ex vivo* duct culture (**Figure 2a**) and found that as ducts become cysts, BECs express ITGA2 protein de novo, which localises to the basal (non-luminal) surface of cysts (**Figure 4c**). Treatment of these *ex vivo* cultures with the integrin-*α*2 *β*1 inhibitor, TCI 15, significantly reduced the area of cystic structures that formed compared to vehicle-treated cultures alone (**Figure 4d**). Moreover, cystic cholangiocytes isolated from patients with PCLD harbouring pathogenic mutations in *GANAB* (c. 2515 C>T) or *PRKCSH* (c. 292+1G>C) treated with TC-I 15 for 48 h demonstrated reduced proliferation (**Figure 4e**), which was not the case for TC-I 15-treated normal human BECs (**Figure 4f**). This growth inhibition was also seen in *Pkhd1^pck^* cystic rat BECs and *Pkhd1^pck^* organoids following TC-115 treatment (**Supplementary Figure 13b** and **13c**), reiterating the requirement of integrin-*α*2*β*1 in the growth of cystic epithelial cells across genetic origins and species.

To test whether integrin-*α*2*β*1 inhibition *in vivo* was sufficient to halt or reverse disease progression in cyst bearing animals, 6-month old *Wdr35^−/−^* mice were dosed with TC-I 15 for 3 weeks (**Figure 4g**). In treated animals, the level of pMLC2 staining in cystic BECs was significantly decreased compared to untreated cysts, indicating that integrin-*α2β*1 signalling *in vivo* directly underlies the cytoskeletal remodelling observed in cystic epithelial cells (**Figure 4h**). Treatment of cyst-bearing animals with TC-I 15 did not reduce cyst size in these animals (in fact, cysts are moderately larger), however significantly fewer cysts formed following treatment without an appreciable difference in serum biochemistry (**Figure 4i** and **Supplementary figure 13d**), indicating that in polycystic disease cysts undergo active, integrin-*α*2*β*1-dependent fission and this cell autonomous process drives disease progression.

## Discussion

Hepatorenal fibrocystic diseases are the most common monogenic diseases in the world(3, 49). The genes that, when mutated, give rise to these conditions are well defined but heterogeneous and converge on the form and function of primary cilia. Despite this, our understanding of the cellular processes that drive cystogenesis particularly in the liver is lacking and as such, we have struggled to define unifying mechanisms by which hepatic cystogenesis can arise. Our data shows that this via TGF*β*-SMAD signalling cystic BECs condition their microenvironment thereby promoting integrin-dependent cyst-fission to generate polycystic disease in a process reminiscent of intestinal crypt fission, where inflation and collapse result in new crypt structures commonly found in pre-neoplastic colonic disease(50–52). We suggest that cyst-fission in PCLD is unlikely a passive process of inflation and collapse, rather septa driven by reactivation of tissue morphogenesis and cytoskeletal remodelling actively drives this process and we propose is likely to generate cysts in liver pathologies in which groups of cysts remain interlinked and connected to the biliary tree such as CHF and autosomal recessive polycystic liver disease. Cyst fission reveals new pharmacological targets that can be leveraged to limit the formation of cysts across multiple tissues.

## Supporting information

Supplemental Materials, Methods and Acknowledgements

## Supplementary tables

- **Supplementary table 1:** Differentially expressed genes between cluster 1 and 2 following single cell analysis.
- **Supplementary table 2:** Enriched GO and KEGG terms when comparing scRNA from cluster 1 and 2.
- **Supplementary table 3:** Differentially expressed genes between vehicle and SIS3 treated cyst-bearing animals.
- **Supplementary table 4:** GOrilla and REViGO outputs from DEGs presented in **Supplementary table 3**.
- **Supplementary table 5:** Putative cell autonomous ligand-receptor interactions in wild-type epithelial cells.
- **Supplementary table 6:** Putative cell autonomous ligand-receptor interactions in cystic epithelial cells.
- **Materials table 1:** Clinical characteristics of human tissue samples.
- **Materials table 2:** Antibodies used in this study.

---

**Supplementary Figure 1:**
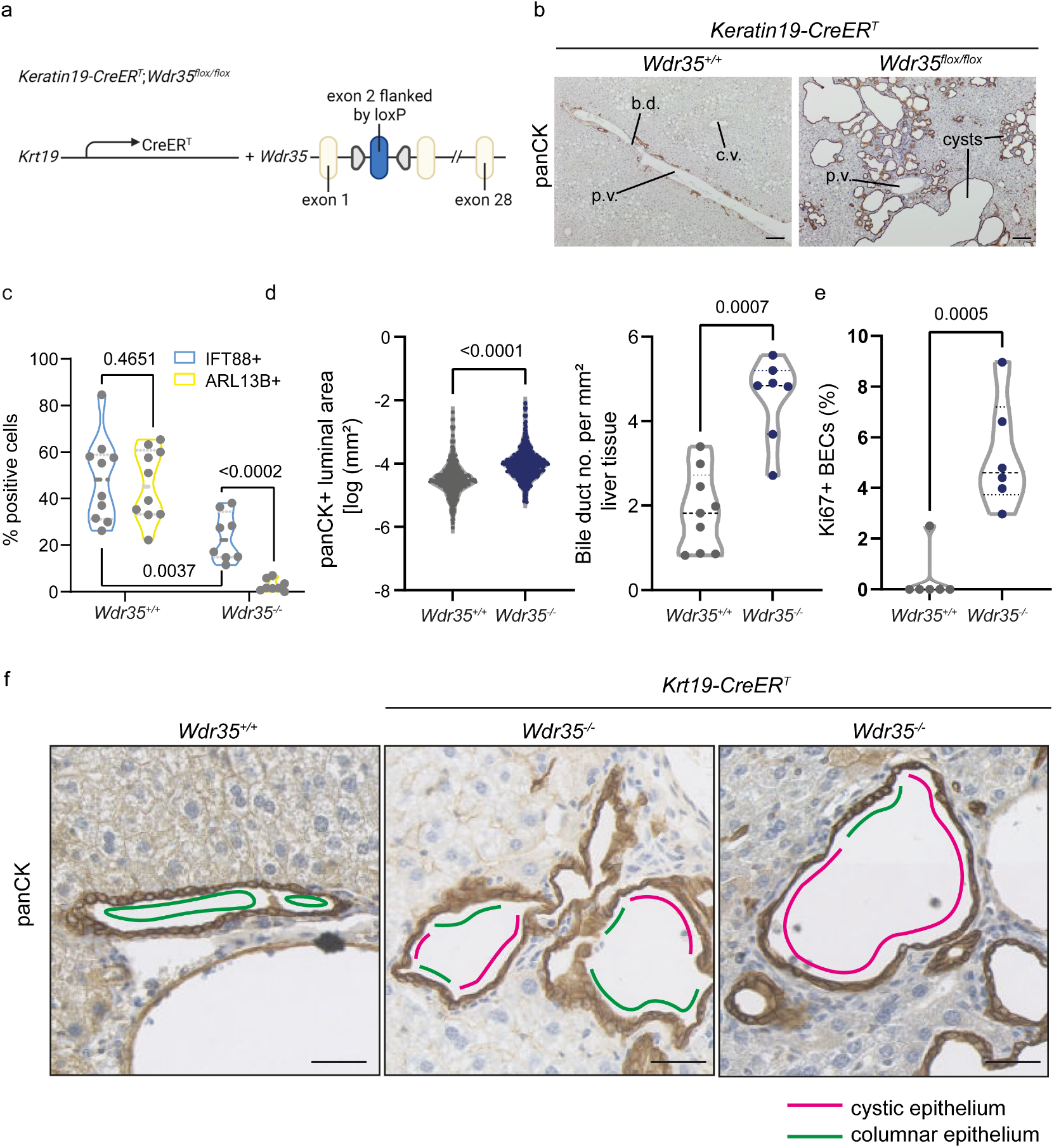
Loss of *Wdr35* in BECs promotes increased cyst formation over time. **a.** Schematic of the genetic strategy for deleting Wdr35 in biliary cells. **b.** Immunohistochemistry of *Wdr35^+/+^* and *Wdr35^−/−^* livers stained for panCK (brown). b.d.-bile duct, c.v.-central vein, p.v.-portal vein **c.** Quantification of IFT88 and ARL13B positive cilia in *Wdr35^+/+^* (N=10 mice) and *Wdr35^−/−^* BECs (N=8 mice). **d.** Luminal area of cysts (*Wdr35^+/+^* n=426 ducts from 7 mice and *Wdr35^−/−^* n=659 cysts from 6 mice), left panel, and cyst number per mouse (*Wdr35^+/+^* N=7 and *Wdr35^−/−^* N=9), right panel, six months following Wdr35-deletion.**e.** Proportion of Ki67 positive (proliferative) cells in cysts 12 months following Wdr35-loss (*Wdr35^+/+^* N=6 animals and *Wdr35^−/−^* N=6 animals). **f.** Immunohistochemistry for panCK on *Wdr35^+/+^* and *Wdr35^−/−^* at 6 months (middle panel) and 12 months (right panel). Green lines denote duct regions where cells are columnar and magenta lines are regions occupied by cystic epithelial cells. Scale bar=100 *μ*m.

**Supplementary Figure 2:**
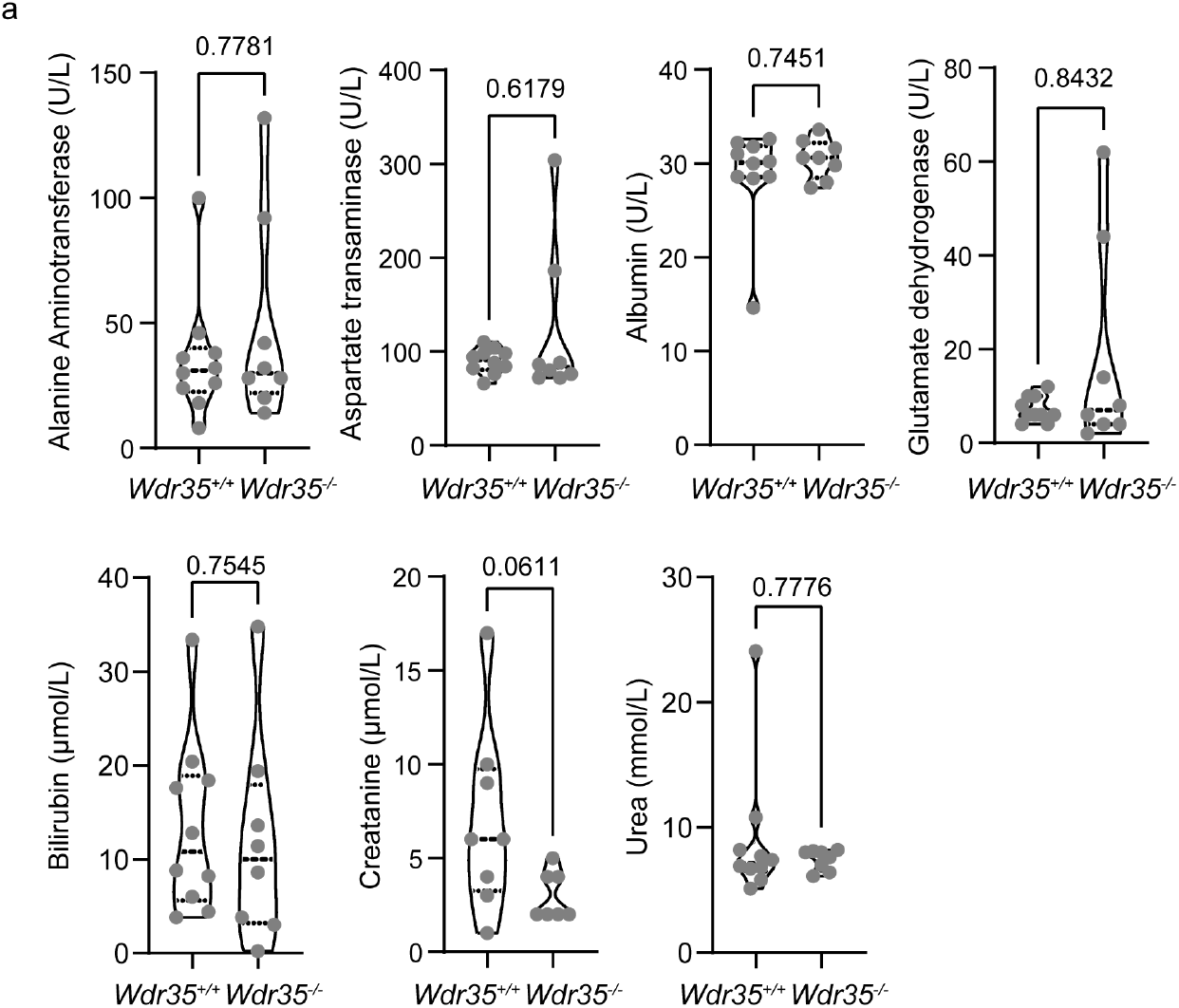
The loss of cilia in murine BECS in adult animals does not universally affect liver function. **a.** Blood serum biochemistry from animals in which *Wdr35* is deleted in BECs 12 months following deletion *Wdr35^+/+^* (N=10 mice) and *Wdr35^−/−^* BECs (N=8 mice). See also **Figure 1d**.

**Supplementary Figure 3:**
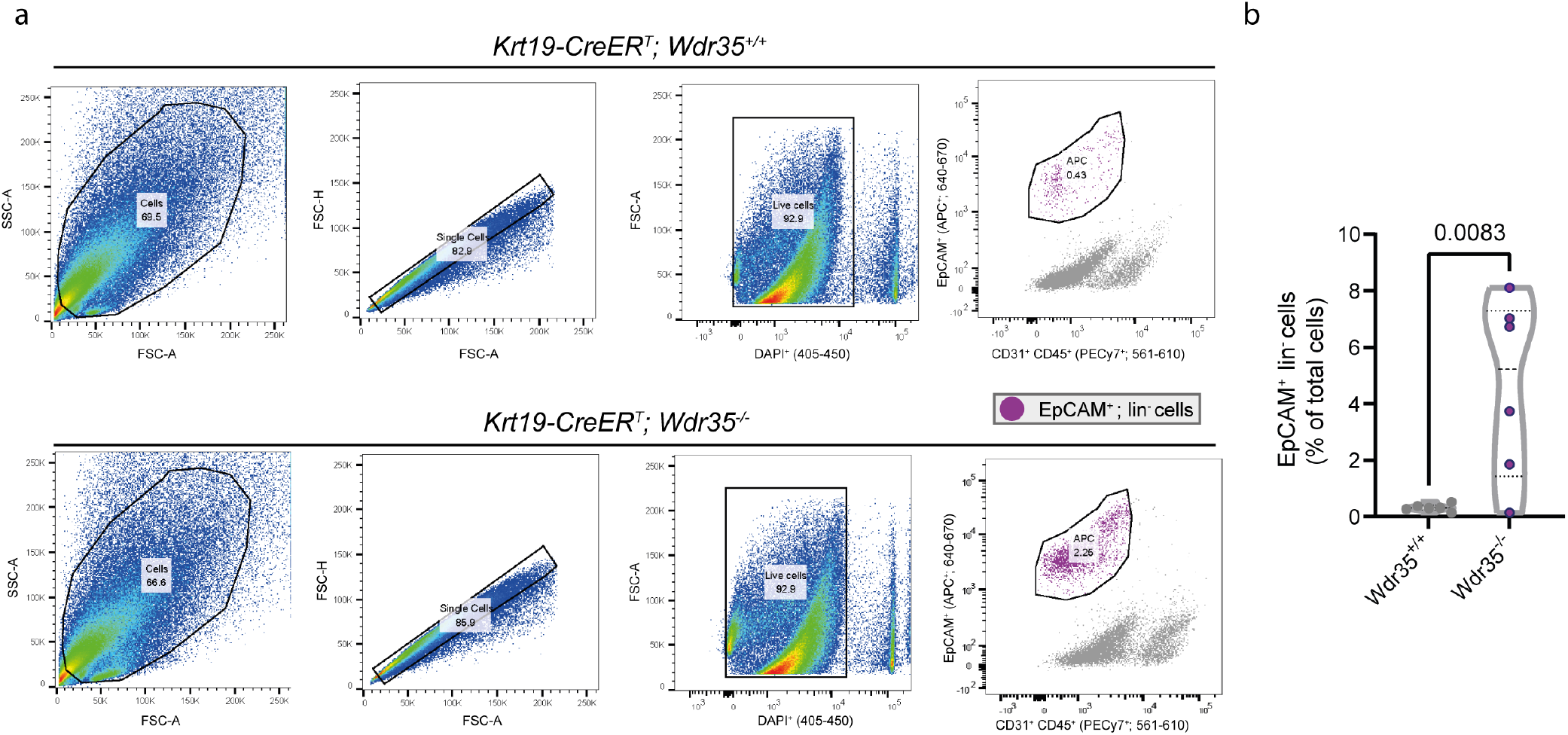
Normal and cystic BECs can be isolated from the liver using EpCAM. **a.** Flow cytometry gating strategy for single cell sequencing. Briefly, single cells were stained with DAPI. Live cells were then gated for CD45- and CD31- and EpCAM+. CD45-/CD31-/EpCAM+ cells were used as input for scRNAseq. **b.** Proportion of total cells that are positive for EpCAM (N=6 mice per genotype).

**Supplementary Figure 4:**
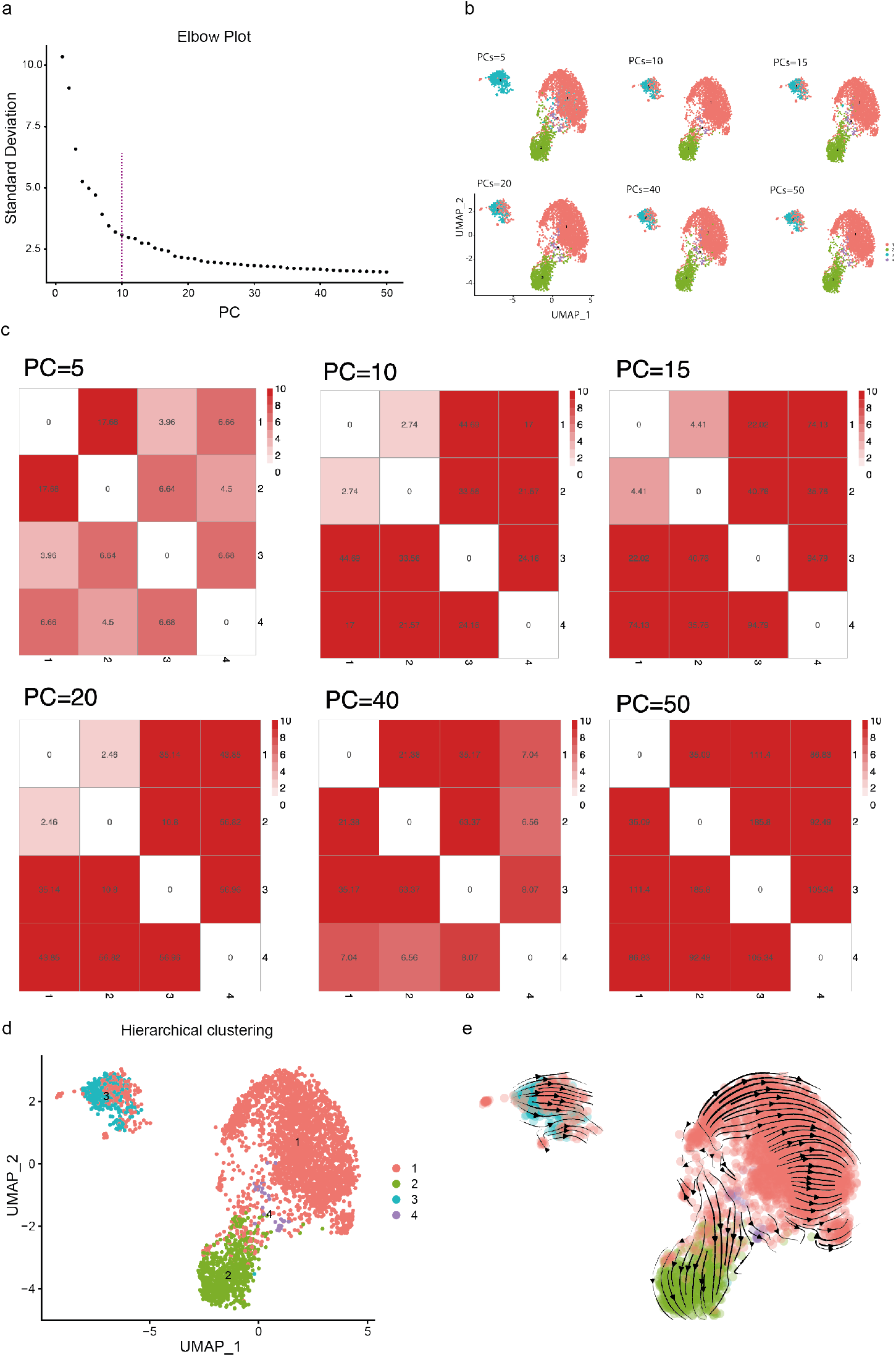
Cystic BECs are transcriptionally distinct from normal BECs. **a.** The Elbow method was used to determine the optimal number of clusters within the scRNA dataset. The dashed line at 10 Principal Components (PCs) corresponds with the point of inflection. **b.** Ward-linkage hierarchical clustering with squared Euclidean distance was used to obtain 4 clusters and determine cluster stability across different PCs. Increasing from PC5 to PC10 shows a slight difference however beyond PC10 clustering remains stable. **c.** Heatmap showing the log_1_ 0 FDR from test difference in mean for all pairs of cluster. The difference between any pairwise clusters is significant. **d.** PC10 was determined as the appropriate input for subsequent analyses. **e.** UMAP embedding of BECs, coloured by identified clusters and overlaid with the RNA velocity stream.

**Supplementary Figure 5:**
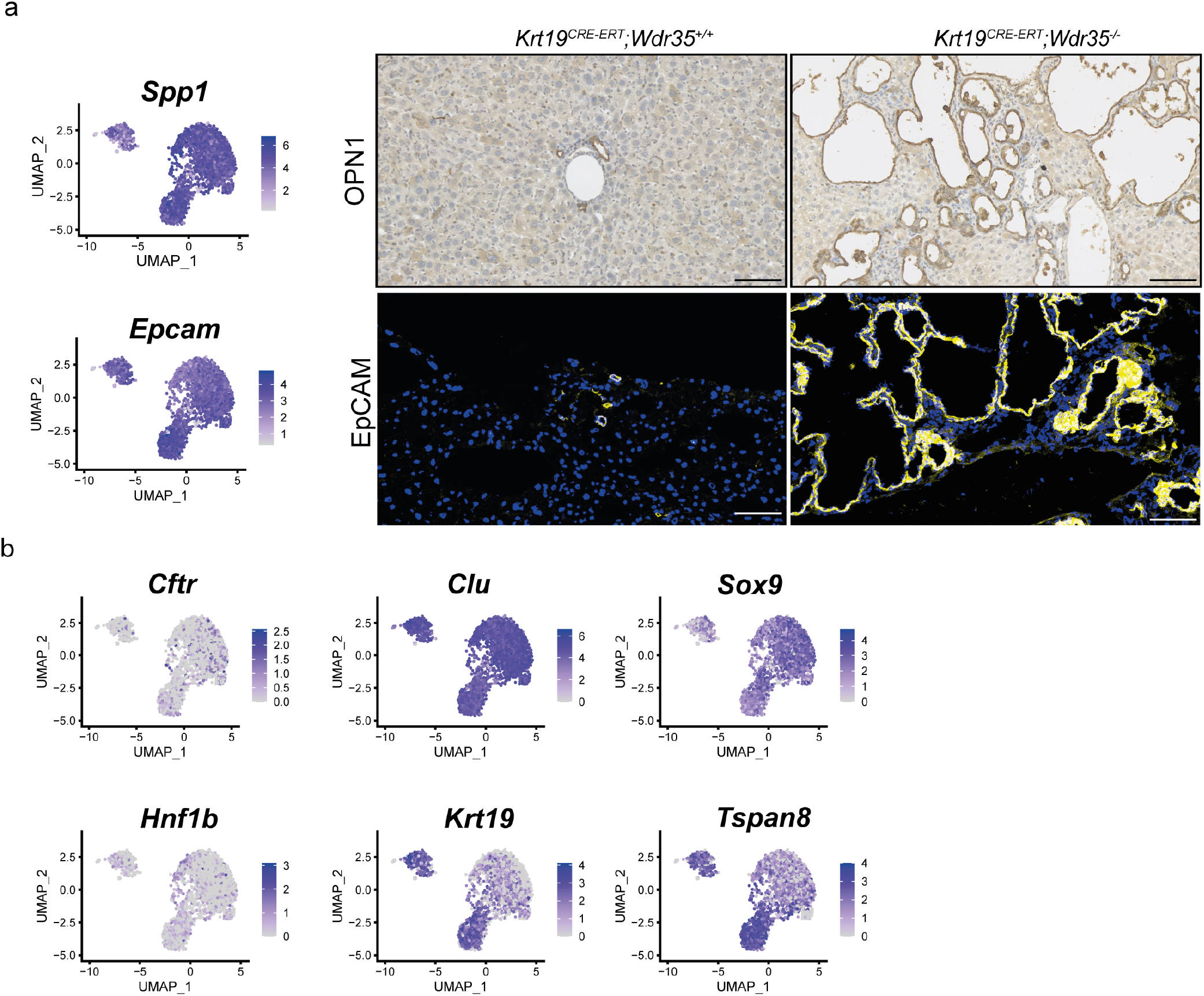
BEC markers are consistently expressed across normal and cystic cells. **a.** Transcript expression of *Spp1* and *Epcam* in both *Wdr35^+/+^* and *Wdr35^−/−^* BECs. Immunohistochemistry showing immunopositivity of OPN1 and EpCAM (yellow) in both normal and cystic BECs (scale bar=100 *μ*m). **b.** Transcript expression of BEC genes *Cftr*, *Clu*, *Sox9*, *Hnf1β*, *Krt19* and *Tspan8.*

**Supplementary Figure 6:**
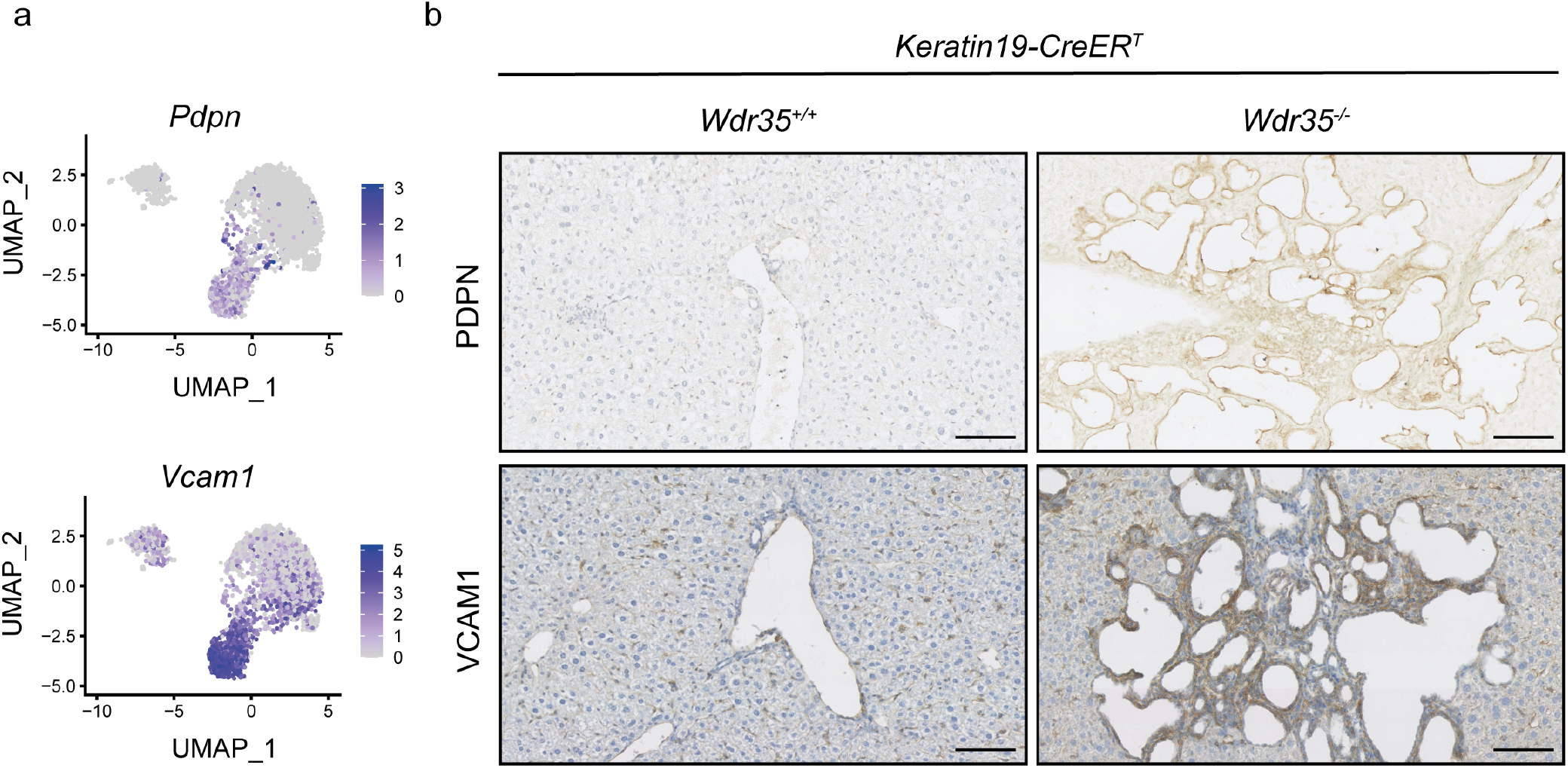
Cystic BECs express specific subsets of genes. **a.** Transcript expression of *Pdpn* and *Vcam1* specifically in cystic epithelial cells. **b.** Immunohistochemistry for PDPN and VCAM1 in *Wdr35^+/+^* BECs and *Wdr35^−/−^* BECs (scale bar=100 *μ*m).

**Supplementary Figure 7:**
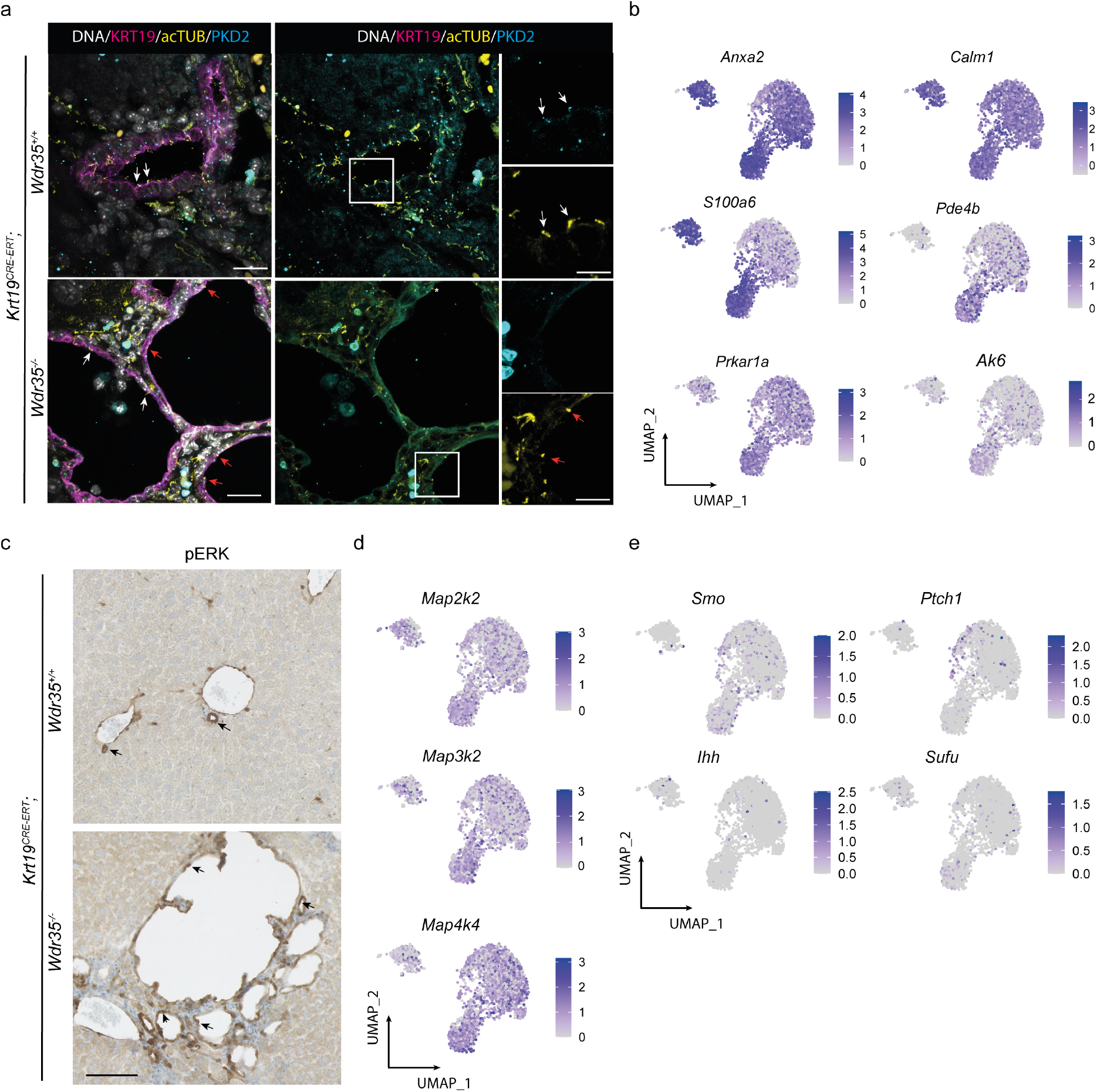
Cystic BECs are enriched for components of Ca^2+^ and MAPK signalling. **a.** Immunofluorescent staining of *Wdr35^+/+^* and *Wdr35^−/−^* ducts stained for KRT19 (magenta), acTUB (yellow), PKD2 (cyan). Scale bar=100 *μ*m, inset scale bar 20 *μ*m. White arrows denote PKD2 (Polycystin-2) expression in acTUB-positive cilia, Orange arrows denote acTUB rudiments lacking PKD2 expression. White box denotes region of magnification. **b**. Transcript expression of Ca^2+^ signalling/PKA components, *Anxa2*, *Calm1*, *S100a6*, *Prkar1a*, *Pde4b* and *Ak6* is enriched in cystic BECs. **c**. pERK immunohistochemistry *Wdr35^+/+^* and *Wdr35^−/−^* ducts arrows denote positive cells. Scale bar=100 *μ*m). **d**. Transcript expression of MAPK pathway intermediates *Map2k2*, *Map3k2* and *Map4k4* are enriched in cystic BECs. **e.** mRNA expression from scRNA of Hedgehog pathway components *Smo*, *Ptch1*, *Ihh*, and *Sufu*.

**Supplementary Figure 8:**
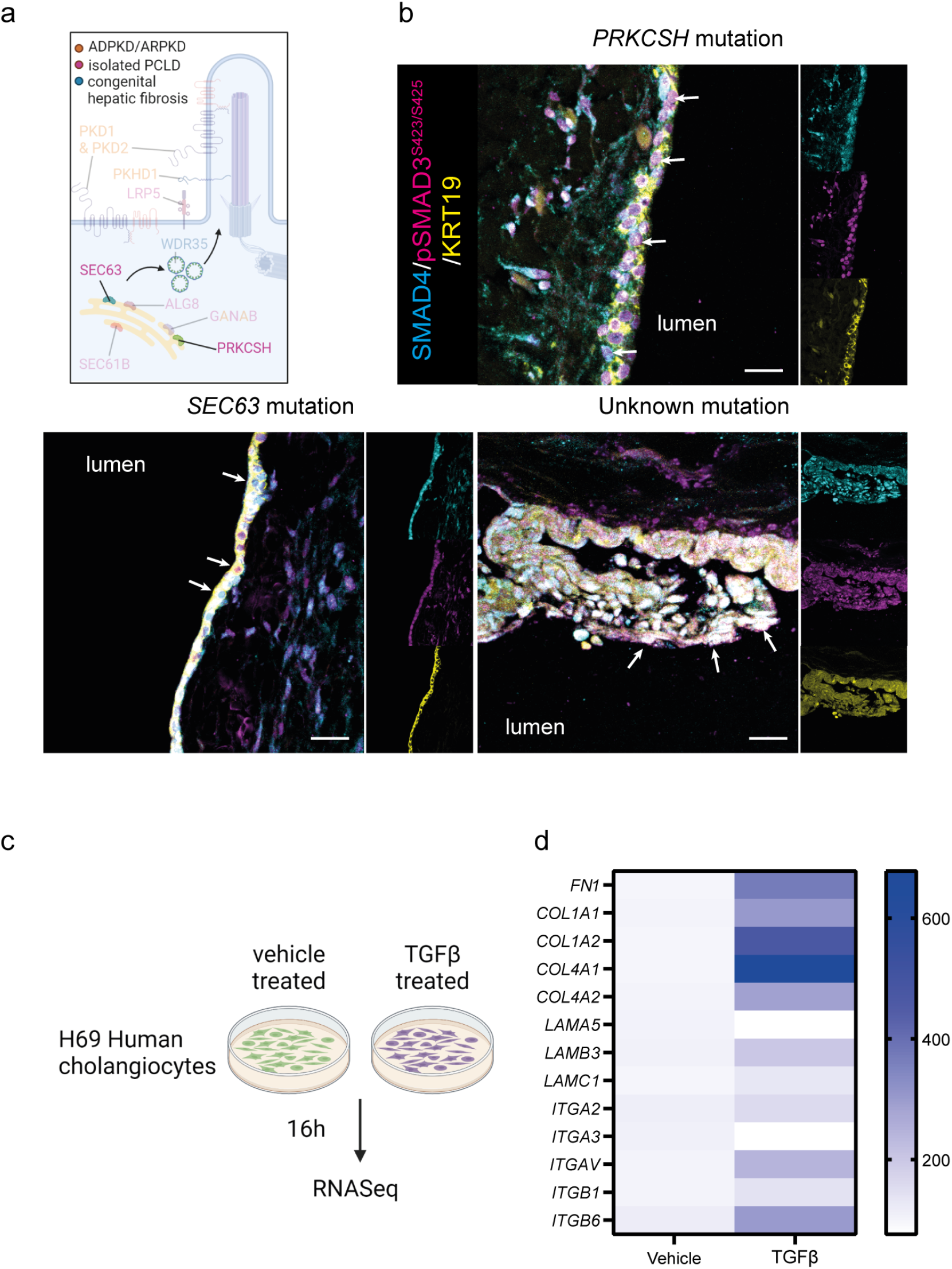
Human BECs treated with recombinant TGŖ*β* express higher levels of ECM molecules. **a.** Diagram demonstrating where isolated polycystic liver disease mutations are within cilia biogenesis. **b.** Immunohistochemistry of SMAD4 (cyan) pSMAD3^*S*423^*1^S^*^425^ (magenta, arrows) KRT19 (yellow) in human cysts with a *PRKCSH, SEC63* mutation or tissue where the causative mutation is unknown. Scale bar=100 *μ*m. **c.** Schematic demonstrating the experimental approach where H69 human cholangiocytes were stimulated with 10 ng/ml TGF*β* for 16 h. **d.** Differential gene expression of ECM genes and integrins in human BECs following TGF*β* stimulation.

**Supplementary Figure 9:**
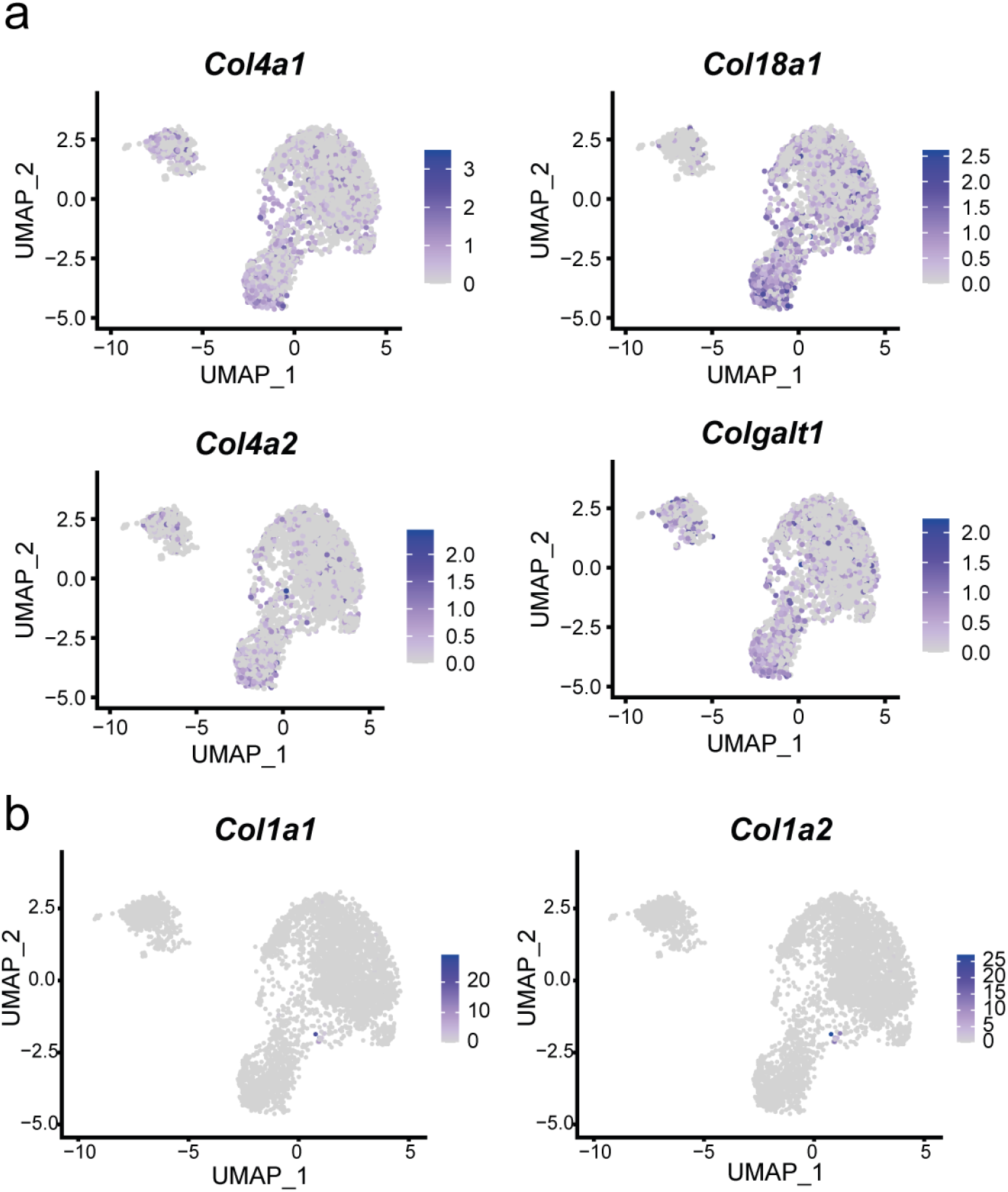
Mouse cystic BECs are enriched for extracellular matrix transcripts and exist in an expanded ECM. **a.** UMAP showing expression of *Col4a1*, *Col4a2*, *Col18a1*, and *Colgalt1.* **b.** UMAP showing expression of *Col1a1* and *Col1a2.*

**Supplementary Figure 10:**
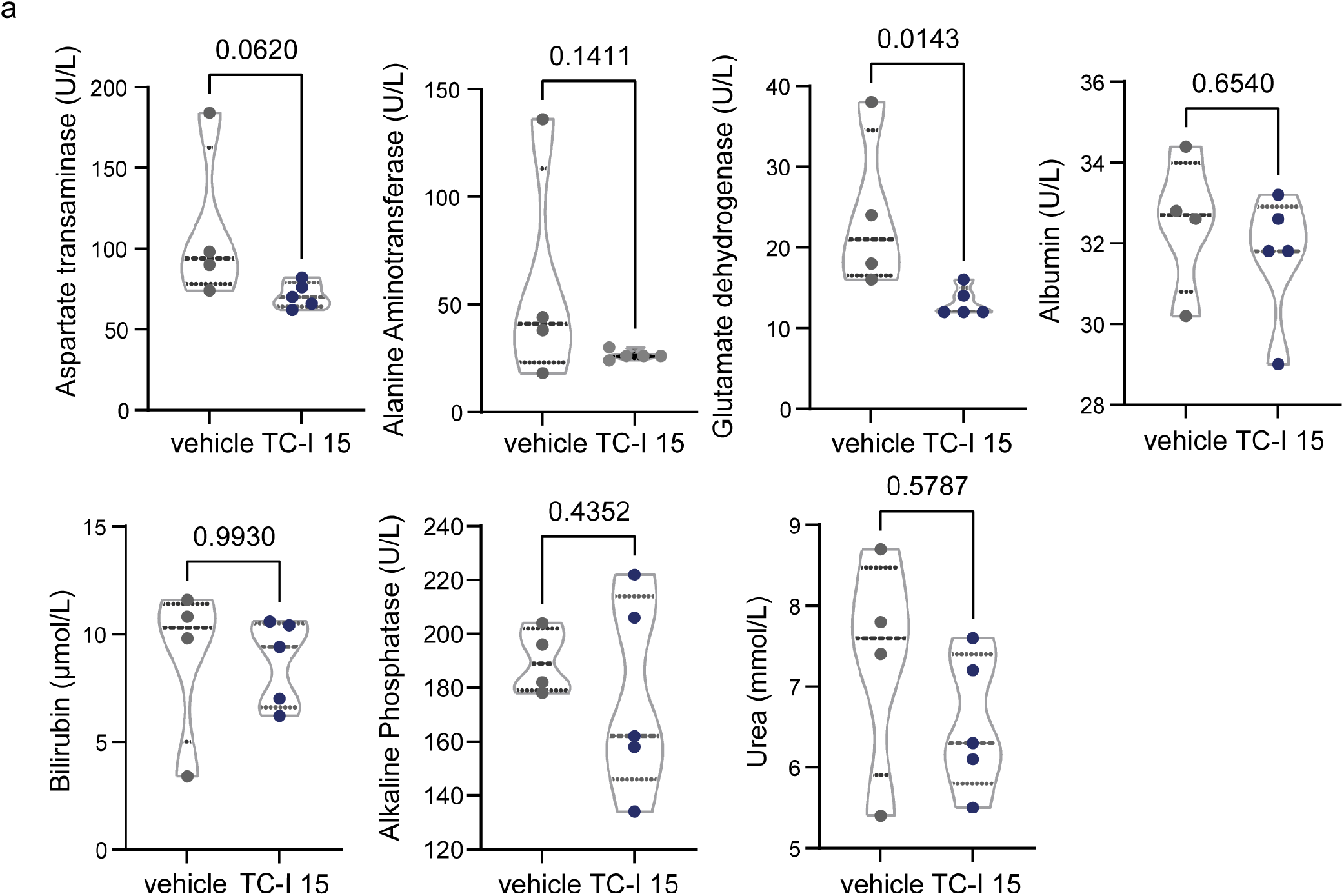
Blood biochemistry from mice treated with the SMAD3 inhibitor, SIS3. **a.** Blood serum biochemistry in *Wdr35^−/−^* cyst baring mice treated with the SMAD3-inhibitor, SIS3 (N=7) or vehicle (N=4) for 3 weeks.

**Supplementary Figure 11:**
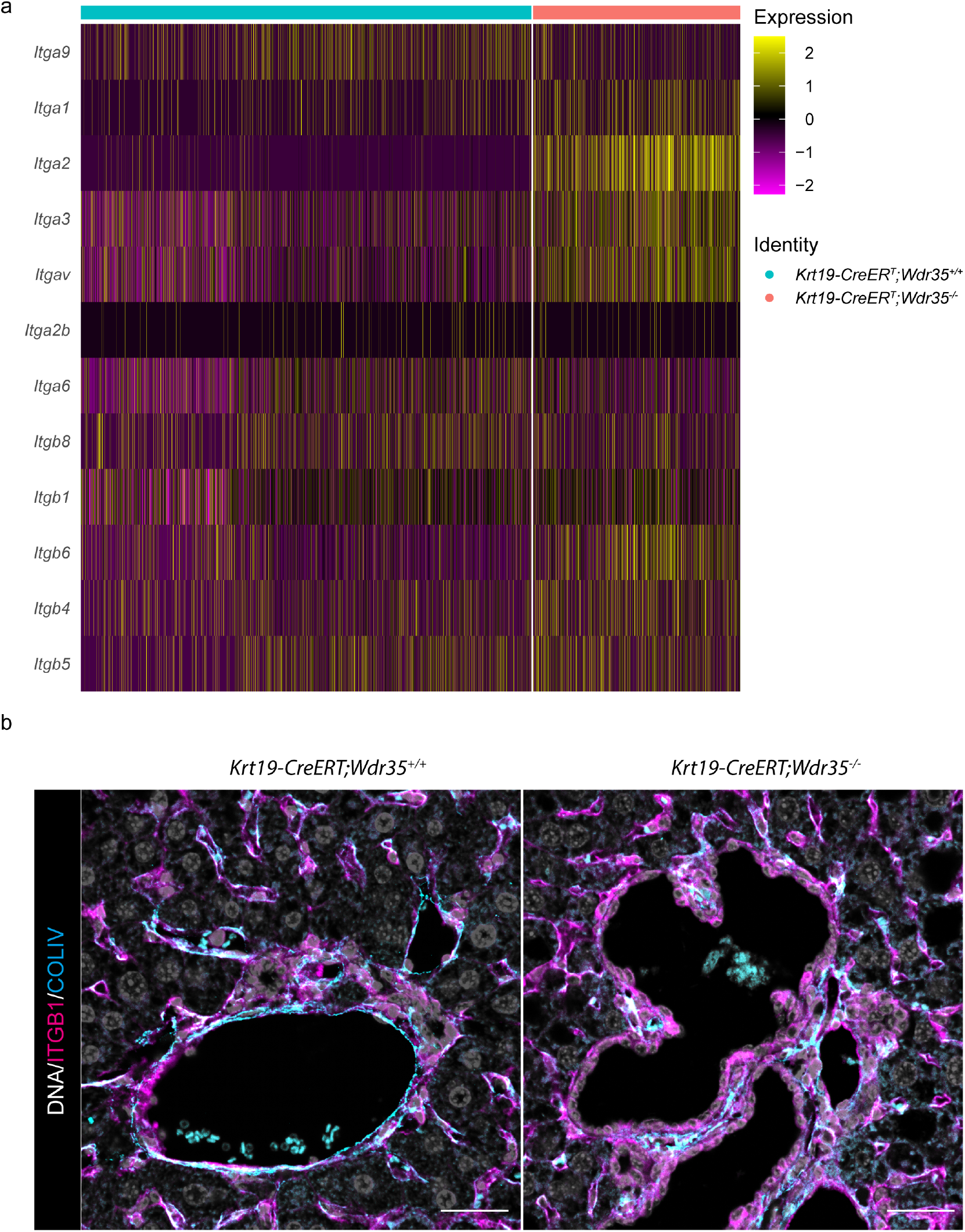
Cystic BECs express a novel profile of integrin molecules. **a.** Heatmap comparing the mRNA expression of all *Itga* and *Itgb* genes that are expressed in single cell RNAseq analysis in BECs from *Wdr35^+/+^* and *Wdr35^−/−^* mice. **b.** Immunofluorescent staining of *Wdr35^+/+^* and *Wdr35^−/−^* livers for ITGB1 (magenta) and COLIV (cyan). DNA is in grey. Scale bar=100 *μ*m.

**Supplementary Figure 12:**
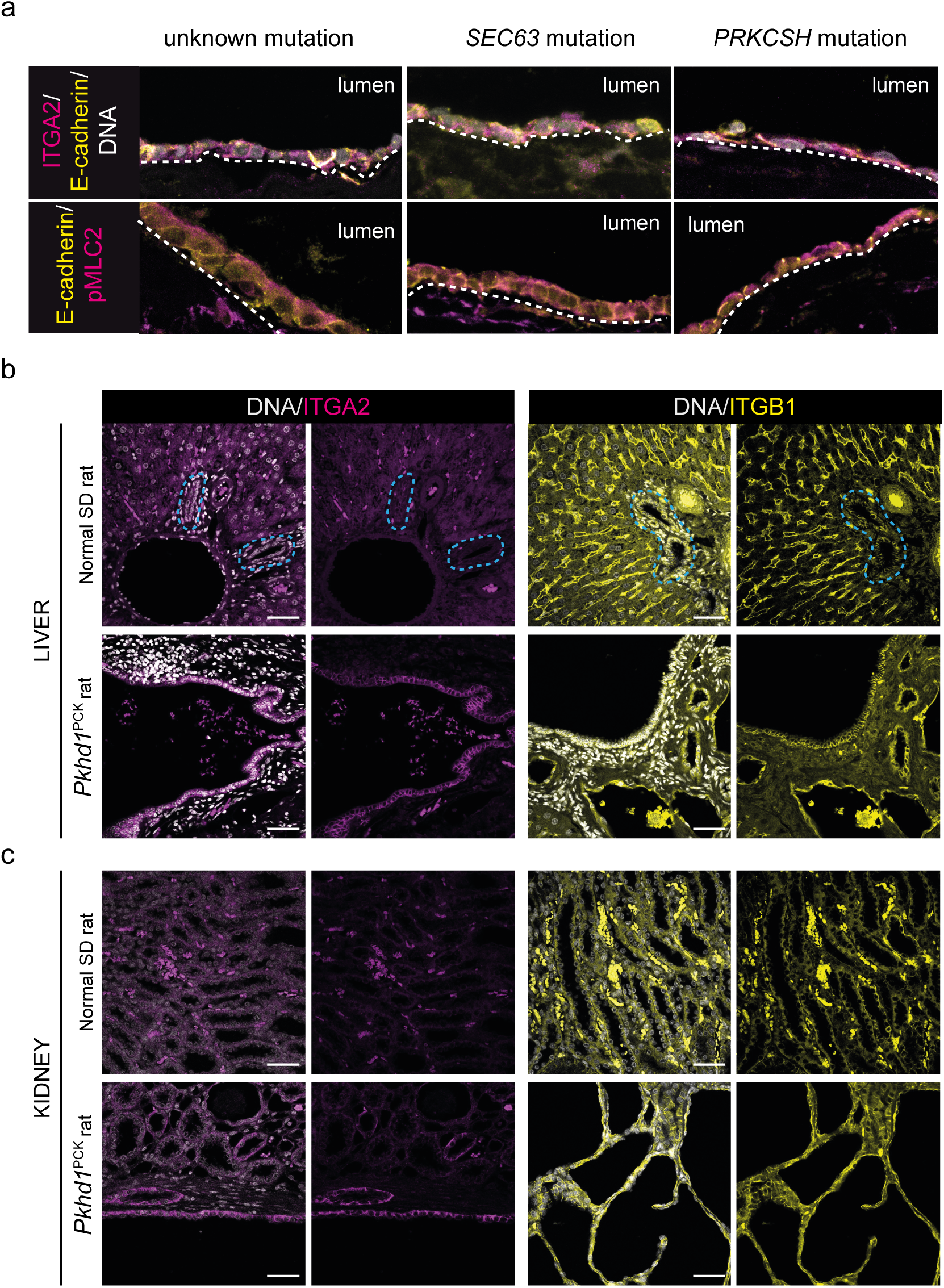
The formation of a pro-cystic state driven by *α2β* 1-integrin is common across tissues and species. **a.** Immunofluorescent staining of human liver cysts with unknown, *SEC63* and *PRKCSH* mutations for DNA (grey), ITGA2 (magenta) and E-cadherin (yellow), upper panels and pMLC2 (magenta) and E-cadherin (yellow), lower panels. **b-c** Immunofluorescent staining of liver (b) and kidney (c) from wild-type rats and rats harbouring a mutation in *Pkdh1* for ITGA2 (magenta) and ITGB1 (yellow). DNA is in grey. Dotted blue line demarcates the bile duct. Scale bar = 100 *μ*m.

**Supplementary Figure 13:**
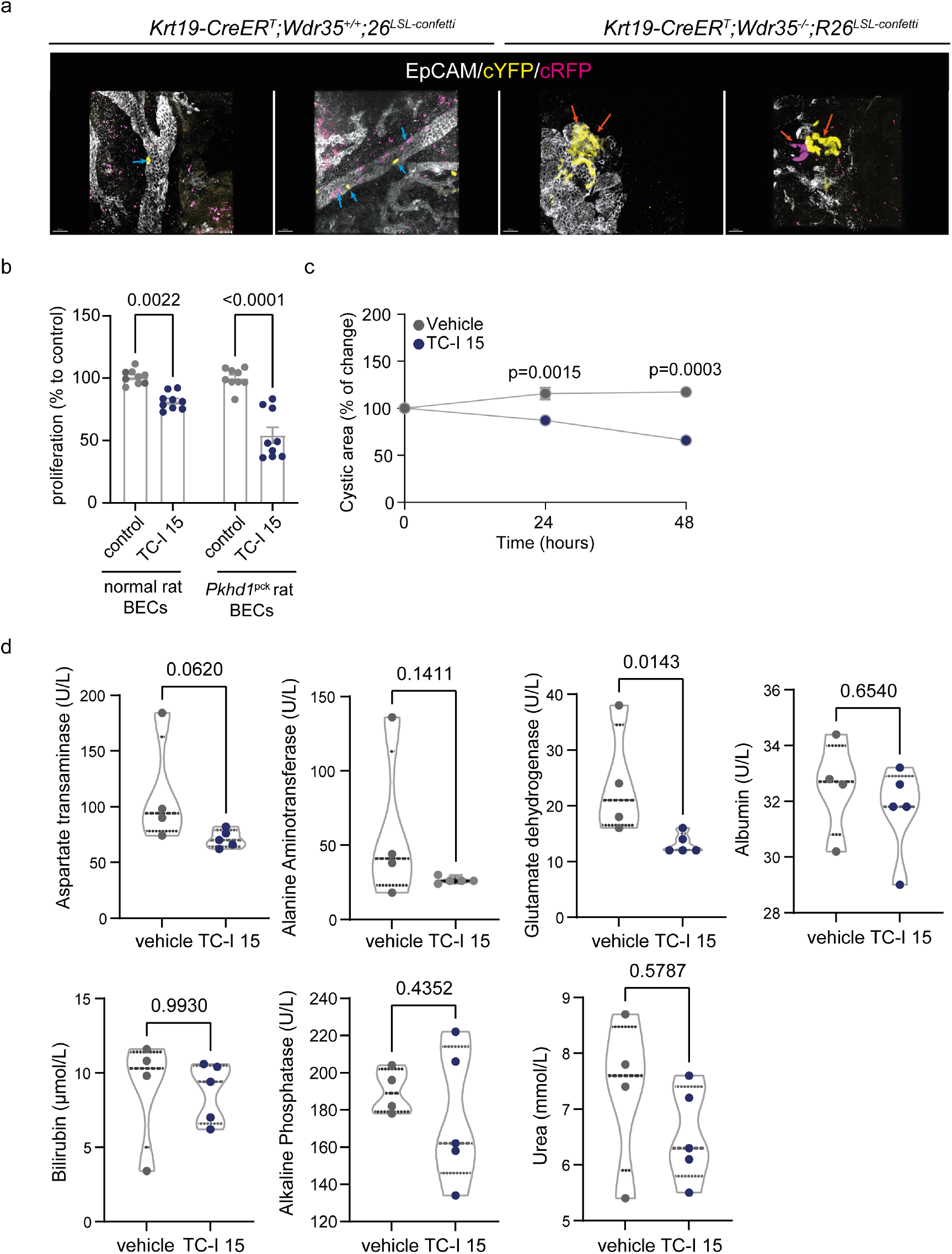
Blood biochemistry from mice treated with the SMAD3 inhibitor, SIS3. **a.** FUnGI cleared wholemount confocal imaging of Confetti fluorophores (cYFP, yellow and cRFP, magenta) in *Wdr35^+/+^* or *Wdr35^−/−^* BECs, 6 months following tamoxifen administration. Blue arrows denote isolated EpCAM positive cells positive for a Confetti fluorophore sporadically along the duct. Orange arrows denote clusters of positive cells forming cysts. **b**. Proliferation rates in normal rat BECs (left hand bars) versus *Pkhd1*-mutant BECs (right-hand bars) (N=9 experimental replicates per group). **c**. Area of cystic rat *Pkdh1*-mutant organoids that form following vehicle versus TC-I 15 treatment (N=3 biological replicates). **d**. Serum biochemistry from cyst bearing *Wdr35^−/−^* mice treated with vehicle alone (N=4 mice) or the integrin *α2β* 1-inhibitor, TC-I 15 (N=5 mice).

## Online Materials and Methods

### Human Tissue

Human normal liver tissue and liver from patients with liver cysts were obtained from the NRS BioResource and Dr Tim Kendall, NHS consultant pathologist, confirmed pathology. JPH Drenth, Department of Gastroenterology and Hepatology, Institute kindly provided samples from patients with PCLD with known genetics for Molecular Life Sciences, Radboud University Medical Center. All tissues were collected under informed consent or were archival diagnostic specimens. Details of human tissue is summarised in **Supplementary table 1**.

### Animal models

#### Transgenic deletion of *Wdr35*

Mice carrying floxed alleles of exon 2 of *Wdr35 Wdr35^flox/flox^*(1) were crossed with animals expressing *Keratin19-CreER^T^* (2). Animals were bred such that all mice contained *CreER^T^*, experimental animals were homozygous for the Wdr35flox/flox allele and controls were wild type (*Wdr35^+/+^). Krt19-CreER^T^;Wdr35^+/+^* or *Krt19-CreER^T^;Wdr35^flox/flox^* animals were treated with tamoxifen (three times 4 mg doses at five weeks old) either by IP injection or oral gavage to activate CRE. Following tamoxifen treatment, transgenic animals that have lost *Wdr35* are denoted as *Wdr35*^−/−^. Animals were housed in same sex groups in 12 h lightdark cycles, with access to food and water ad libitum. Both males and females were used in this study. For experiments containing the *R26*^LSL–Confetti/+^ reporter(3), *Krt19-CreER^T^;Wdr35^flox/flox^* were crossed to generate *K19CreER^T^*;*Wdr35^flox/flox^*;*R26^LSL–Confetti/+^* mice and were treated with tamoxifen as detailed above. For inhibitor studies, cyst bearing animals were treated for 3 weeks with 20 mg/kg of TC-I 15 (a selective integrin *α2β1* inhibitor), 10 mg/kg of SIS3 (an inhibitor of SMAD3) or vehicle alone. Animals were maintained in SPF environment and studies carried out in accordance with the guidance issued by the Medical Research Council in “Responsibility in the Use of Animals in Medical Research” (July 1993) and licensed by the Home Office under the Animals (Scientific Procedures) Act 1986. Experiments were performed under project license number PFD31D3D4 in facilities at the University of Edinburgh (PEL 60/6025).

#### Blood biochemistry

At necropsy, blood was collected from the inferior vena cava and spun at 12000 g to pellet the cellular fraction. Serum was collected and provided to the SURF facility, Queens Medical Research Institute, Edinburgh where they performed ELISA for aspartate transaminase, alanine Aminotransferase, glutamate dehydrogenase, albumin, bilirubin and urea.

#### Fluorescent activated cell sorting and 10X single cell sequencing

12 month *Krt19-CreER^T^;Wdr35^flox/flox^* or *Krt19-CreER^T^;Wdr35^+/+^* mice were perfused with saline and livers were digested with collagenase and dispase to enrich for the biliary tree. Enriched bile ducts were dissociated into single cells using trypsin and stained for EpCAM-APC, CD31-PECy7, CD45-PECy7 (**Supplementary table 2**). Live cells (which are negative for DAPI) were identified and CD31-PECy7-/CD45-PECy7-/EpCAM+ cells were isolated and 10,000 cells were used as input for Chromium Next GEM Single Cell 3’ analysis by the FACS and Single Cell Core at the MRC Human Genetics Unit Edinburgh. Generated libraries were quantified and sequenced on an Illumina P1000 flow cell by the Clinical Research Facility, University of Edinburgh. All RNA sequencing data pertaining to this manuscript is deposited on the NCBI Gene Expression Omnibus (GEO) as accession number GSExxxxxx. Bulk RNA data from TGF*β*-treated H69 cells was downloaded from GSE145127.

#### Analysis of single cell RNAseq data

Raw sequencing data was analyzed using Cell Ranger’s pipelines (version 5.0.0) provided by 10x Genomics. Specifically, ‘cellranger mk-fastq’ was initially used to generate FASTQ files from raw base call files. Then, ‘cellranger count’ was applied to align FASTQ files to the pre-build mouse reference mouse genome (GRCh38) provided by Cell Ranger and generate single cell feature counts.

Seurat (v4.0.6, https://satijalab.org/seurat/) was used to filter out cells: (1) with unique feature counts over 7,000 or less than 600, (2) with counts greater than 50,000, (3) with more than 5% mitochondrial counts, (4) without expression of BEC marker *Epcam* and *Sppl.* DoubletFinder (v2.0) was used to predict and remove technical artifacts or ‘doublets’ for each sample. Specifically, estimated doublet rates were calculated according to the 10x Chromium User Guide. Homotypic doublet rates were also taken into account using the modelHomotypic function with default parameters. We subsequently removed genes that are present in less than 3 cells in each sample.

After the preprocessing and quality filtering steps, the dataset was then processed using the standard Seurat workflow: 1. log normalize data using the NormalizeData function, with a default size factor 10,000, 2. scale data using the ScaleData function, 3. identify variable features using FindVariableFeatures with “vst” parameters and then imputed into PCA using RunPCA function. RunUMAP function was used for UMAP embedding visualisation with the first 10 PCs (principal components).

Ward-linkage hierarchical clustering was performed with squared Euclidean distance on the first 10 PCs to obtain clusters (arXiv:2012.02936v2 [stat.ME]). Before proceeding to downstream analyses, such as differential gene expression analysis, it is important to avoid the problem of selective inference. Selective inference is described as the assessment of significance as well as effect sizes from the same dataset which has been carried out via statistical tests to find potential associations. Selective inference has recently been demonstrated to have an important role in accurately estimating significance in biomedical science –omic data(4). Specifically, in the scRNA-seq clustering analysis, one first clusters the data, then measures if they are significantly different and identifies differentially expressed genes between clusters. This leads to inflated Type I error rate (artificially deflated p-values). We therefore applied selective inference correction implemented by clusterpval R (arXiv:2012.02936v2 [stat.ME]) package to determine an appropriate number of clusters and confirmed that the identified clusters are pairwise significantly different. We also assessed the stability of estimated clusters under different PC projections. We then used FindMarkers function from Seurat to identify differentially expressed genes (DEGs) whose log-fold changes are greater than 0.25, BH-adjusted p-values less than 0.05 by the Wilcox test and are expressed by at least 25% of cells. The clusterProfiler package was used to perform Gene Set Enrichment Analysis for DEGs. The enrichKEGG and enrichGO function were used to identify significant GO (Gene Ontology) terms and KEGG (Kyoto Encyclopedia of Genes and Genomes), respectively, with threshold p value < 0.05 and q value < 0.05. The background gene set of this analysis was selected from publication literature(5), which includes a list of genes that could be expressed in the EpCam sorted cholangiocytes from mouse. R package ggplot2 was used to visualise the results.

We performed cellular trajectory analysis by RNA velocity. We first used velocity(6) to generate the loom files from prealigned bam files. The loom files contain two count matrices of unspliced and spliced RNA abundances.

The proportions of spliced and unspliced counts are 80% and 20% respectively, which is in the appropriate range of velocity inference(7). We then calculated the RNA velocity from scVelo(7) using deterministic (steady-state), stochastic and dynamical mode. They produced consistent results.

Receptor-ligand analysis was performed using R package SingleCellSingalR(8). Gene expression data and hierarchical clustering were used as input to compute receptor-ligand interactions scores between clusters using the ‘cell_signaling’ function. We filtered out receptor-ligand interactions whose LRscores were lower than 0.5. Receptor-ligand interactions were visualized by the ‘visualize_interactions’ function.

#### Isolation of RNA and DNA

RNA was extracted from FACS isolated cells and extracted by TRIzol RNA Isolation Reagent (Invitrogen) lysis. RNA was precipitated with chloroform and cleaned up using the RNeasy Mini Kit (QIAGEN) as per the manufacturer’s instructions. For downstream sequencing applications RNA quality (RIN score) was quantified using the Agilent 2100 Bioanalyzer with an RNA 6000 chip. A minimum RIN threshold of 8 was used for RNA-seq.

#### RNA sequencing data processing and analysis

The primary RNA-Seq processing, quality control to transcriptlevel quantitation, was carried out using nf-core/rnaseq v1.4.3dev (https://github.com/ameynert/rnaseq)(9). Reads were mapped to the mouse FVB_NJ1 decoy-aware transcriptome using the salmon aligner (1.1.0). RNA-Seq analysis was performed in R (4.0.2), Reads were summarized to genelevel and differential expression analysis was performed using the bioconductor packages tximport (1.16.1) and DESeq2 (1.28.1). A pre-filtering was applied to keep only genes that have at least 10 reads in a group and 15 reads in total. The Wald test was used for hypothesis testing for pairwise group analysis. A shrunken log2 fold changes (LFC) was also computed for each comparison using the adaptive shrinkage estimator from the ‘ashr’ package.

#### Duct isolation and culture

Livers were minced finely using razor blades and digested with 0.125 mg/ml collagenase type IV (Gibco) and 0.125 mg/ml dispase II (Gibco) at 37 °C. Once bile ducts were visible they were used for FACS, immunostaining or organoid cultures. For cultures, Isolated bile ducts were suspended in Matrigel (Corning) and cultured in media containing Advanced DMEM/F-12 media (Gibco) containing 1xGlutaMAX (Gibco), 1xAntibiotic-Antimycotic (Gibco), 10 *μM* HEPES (Sigma), 50 ng/ml EGF (RD Systems), 100 ng/ml FGF10 (Novus Biologicals), 5 ng/ml HGF (Novus Biologicals), 10 nM gastrin (Sigma), 10 *μ*M nicotinamide (Acros Organics), 1.25 mM N-acetyl-Lcysteine (Sigma), 1x B27 (Life Technologies), 1x N2 Supplement (Life Technologies), 1 *μ*g/ml R-Spondin-1 (RD Systems), 0.2 *μ*g/ml WNT5A (RD Systems) and 10 nM forskolin (Tocris). Cysts were allowed to culture for 72 hours at 37 °C in a humidified incubator with 5% CO2 before fixing for immunofluorescent staining or lysis for proteomic studies. Inhibition studies had media supplemented with 100 *μ*M TC-I 15 (*α*2*β*1-integrin selective inhibitor), 10 mM SIS3 (SMAD3 selective inhibitor) or DMSO (vehicle control) at equivalent volumes. Media containing inhibitors/vehicle was replaced after 48 hours. Cyst size was determined by brightfield microscopy and measurements made on Fiji (ImageJ).

#### Cell culture

Human (normal human cholangiocytes [NHC], ADPLD and ADPKD) and rat (normal rat cholangio-cytes [NRC] and PCK) cholangiocyte cells were seeded on collagen-coated flasks and cultured in supplemented(10) DMEM/F-12 medium as previously described11. Cell proliferation rates were determined by flow-cytometry using Cell-Trace^™^ CFSE Cell Proliferation Kit (Invitrogen), following the manufacturer’s instructions. Rat 3D cultures were derived from micro-dissected cysts of PCK rats as previously described(11). Isolated tissues were sandwiched between 1.5 mg/ml type I rat tail collagen (BD Biosciences) and cultured in supplemented DMEM/F-12 medium.

#### Immunohistochemistry and quantification

Dissected tissues were fixed overnight in formalin at 4 °C, embedded in paraffin and were sectioned at 4 *μ*m. Following antigen retrieval (see **Supplementary table 2**), tissue sections were incubated with antibodies as detailed in **Supplementary table 2**. Fluorescently stained tissues were counterstained with DAPI prior to imaging. Colorimetric stains were counterstained with Haematoxylin and mounted with DPX. *K19CreER^T^;Wdr35^flox/flox^;R26^LSL–Confetti/+^* and *K19CreER^T^;Wdr35^+/+^;R26^LSL–Confetti/+^* samples were sectioned at 200 *μ*m using a Krumdieck Tissue Slicer and fixed for 45 min in formalin and then cleared using FUnGI clearing as previously described(12, 13). Histological tissues were scanned using a Nanozoomer, using a Nikon A1R or Leica Stellaris confocal microscope and were analysed using either FUJI, Imaris or QuPath.

#### Statistical analysis

All experimental groups were analysed for normality using a D’Agostino–Pearson Omnibus test. Groups that were normally distributed were compared with either a two-tailed Student’s t test (for analysis of two groups) or using one-way ANOVA to compare multiple groups, with a post hoc correction for multiple testing. Non-parametric data were analysed using a Wilcoxon–Mann–Whitney U test when comparing two groups or a Kruskall–Wallis test when comparing multiple non-parametric data. Throughout, P<0.05 was considered significant. Data are represented as mean with S.E.M. for parametric data or median with S.D. for non-parametric data.

#### Figures

All figures were assembled with Adobe Illustrator and graphics were created with BioRender.com.

## ACKNOWLEDGEMENTS

We would like to thank the Advanced Imaging Resource at the IGC, Susan Campbell and Lizzie Freyer at the IGC cytometry and single cell core facility, Richard Clarke at the Clinical Research Facility (University of Edinburgh) and Philippe Gautier for bioinformatics support. JMB is supported by Spanish Carlos III Health Institute (ISCIII) [(FIS PI18/01075, PI21/00922, and Miguel Servet Programme CPII19/00008) cofinanced by “Fondo Europeo de Desarrollo Regional” (FEDER)] and CIBERehd (ISCIII); La Caixa Scientific Foundation (HR17-00601); AMMF-The Cholangiocarcinoma Charity (EU/2019/AMMFt/001); PSC Partners US and PSC Supports UK (06119JB); European Union’s Horizon 2020 Research and Innovation Program [grant number 825510, ESCALON]. PO by the Basque Government (PRE_2016_1_0269).AK was supported by the XDF programme from the University of Edinburgh and Medical Research Council (MC_UU_00009/2). PM is funded by an MRC Unit award (MC_UU_00007/14) and ERC consolidator grant (866355). A Cancer Research UK Fellowship (C52499/A27948) funds LB.

## AUTHOR CONTRIBUTIONS

SHW planned and performed experiments, analysed data and drafted and edited the manuscript. YY analysed data and generated figures for the manuscript. PO designed and performed experiments and analysed data. EJ analysed data and EC performed experiments for the manuscript. KG provided technical support and performed the animal work. JPHD provided human samples in this study. TJK provided clinical pathological support and clinical tissue. JB provided intellectual input, tissue and cells. AK provided analytical support, supervised scRNA-seq analysis, and edited the manuscript. PM and LB lead the project, designed and carried out experiments, analysed data, wrote and edited the manuscript and funded the project.

## Bibliography

1. Wybrich R Cnossen and Joost P H Drenth. Polycystic liver disease: an overview of pathogenesis, clinical manifestations and management. Orphanet Journal of Rare Diseases, 9: 69, may 2014. doi: 10.1186/1750-1172-9-69.

2. Matthew B Lanktree, Amirreza Haghighi, Elsa Guiard, Ioan-Andrei Iliuta, Xuewen Song, Peter C Harris, Andrew D Paterson, and York Pei. Prevalence estimates of polycystic kidney and liver disease by population sequencing. Journal of the American Society of Nephrology, 29(10):2593–2600, oct 2018. doi: 10.1681/{ASN}.2018050493.

3. Luca Fabris, Romina Fiorotto, Carlo Spirli, Massimiliano Cadamuro, Valeria Mariotti, Maria J Perugorria, Jesus M Banales, and Mario Strazzabosco. Pathobiology of inherited biliary diseases: a roadmap to understand acquired liver diseases. Nature Reviews. Gastroenterology & Hepatology, 16(8):497–511, aug 2019. doi: 10.1038/s41575-019-0156-4.

4. Anatoliy I Masyuk, Sergio A Gradilone, Jesus M Banales, Bing Q Huang, Tatyana V Masyuk, Seung-Ok Lee, Patrick L Splinter, Angela J Stroope, and Nicholas F Larusso. Cholangiocyte primary cilia are chemosensory organelles that detect biliary nucleotides via P2Y12 puriner-gic receptors. American Journal of Physiology. Gastrointestinal and Liver Physiology, 295 (4):G725–34, oct 2008. doi: 10.1152/ajpgi.90265.2008.

5. Guangrui Geng, Yunming Xiao, Yingjie Zhang, Wanjun Shen, Jiaona Liu, Fei Zhu, Xu Wang, Jie Wu, Ran Liu, Guangyan Cai, Xueyuan Bai, Qinggang Li, and Xiangmei Chen. Ganab haploinsufficiency does not cause polycystic kidney disease or polycystic liver disease in mice. BioMed research international, 2020:7469428, may 2020. doi: 10.1155/2020/7469428.

6. Kurt A Zimmerman, Cheng Jack Song, Nancy Gonzalez-Mize, Zhang Li, and Bradley K Yoder. Primary cilia disruption differentially affects the infiltrating and resident macrophage compartment in the liver. American Journal of Physiology. Gastrointestinal and Liver Physiology, 314(6):G677–G689, jun 2018. doi: 10.1152/ajpgi.00381.2017.

7. Meral Gunay-Aygun, Maya Tuchman, Esperanza Font-Montgomery, Linda Lukose, Hailey Edwards, Angelica Garcia, Surasawadee Ausavarat, Shira G Ziegler, Katie Piwnica-Worms, Joy Bryant, Isa Bernardini, Roxanne Fischer, Marjan Huizing, Lisa Guay-Woodford, and William A Gahl. PKHD1 sequence variations in 78 children and adults with autosomal recessive polycystic kidney disease and congenital hepatic fibrosis. Molecular Genetics and Metabolism, 99(2):160–173, feb 2010. doi: 10.1016/j.ymgme.2009.10.010.

8. Seokho Kim, Hongguang Nie, Vasyl Nesin, Uyen Tran, Patricia Outeda, Chang-Xi Bai, Jacob Keeling, Dipak Maskey, Terry Watnick, Oliver Wessely, and Leonidas Tsiokas. The polycystin complex mediates wnt/ca(2+) signalling. Nature Cell Biology, 18(7):752–764, jul 2016. doi: 10.1038/ncb3363.

9. Shixuan Wang, Jingjing Zhang, Surya M Nauli, Xiaogang Li, Patrick G Starremans, Ying Luo, Kristina A Roberts, and Jing Zhou. Fibrocystin/polyductin, found in the same protein complex with polycystin-2, regulates calcium responses in kidney epithelia. Molecular and Cellular Biology, 27(8):3241–3252, apr 2007. doi: 10.1128/{MCB}.00072-07.

10. Whitney Besse, Ke Dong, Jungmin Choi, Sohan Punia, Sorin V Fedeles, Murim Choi, Anna-Rachel Gallagher, Emily B Huang, Ashima Gulati, James Knight, Shrikant Mane, Esa Tah-vanainen, Pia Tahvanainen, Simone Sanna-Cherchi, Richard P Lifton, Terry Watnick, York P Pei, Vicente E Torres, and Stefan Somlo. Isolated polycystic liver disease genes define effectors of polycystin-1 function. The Journal of Clinical Investigation, 127(5):1772–1785, may 2017. doi: 10.1172/{JCI90129}.

11. Alvaro Santos-Laso, Laura Izquierdo-Sanchez, Pedro M Rodrigues, Bing Q Huang, Mikel Azkargorta, Ainhoa Lapitz, Patricia Munoz-Garrido, Ander Arbelaiz, Francisco J Caballero-Camino, Maite G Fernández-Barrena, Raul Jimenez-Agüero, Josepmaria Argemi, Tomas Aragon, Felix Elortza, Marco Marzioni, Joost P H Drenth, Nicholas F LaRusso, Luis Bu-janda, Maria J Perugorria, and Jesus M Banales. Proteostasis disturbances and endoplasmic reticulum stress contribute to polycystic liver disease: New therapeutic targets. Liver International, 40(7):1670–1685, jul 2020. doi: 10.1111/liv.14485.

12. Carlos A Bacino, Shweta U Dhar, Nicola Brunetti-Pierri, Brendan Lee, and Penelope E Bon-nen. WDR35 mutation in siblings with sensenbrenner syndrome: a ciliopathy with variable phenotype. American Journal of Medical Genetics. PartA. 158A(11):2917–2924, nov 2012. doi: 10.1002/ajmg.a.35608.

13. Ranad Shaheen, Saud Alsahli, Nour Ewida, Fatema Alzahrani, Hanan E Shamseldin, Nisha Patel, Awad Al Qahtani, Homoud Alhebbi, Amal Alhashem, Tarfa Al-Sheddi, Rana Alomar, Eman Alobeid, Mohamed Abouelhoda, Dorota Monies, Abdulrahman Al-Hussaini, Muneerah A Alzouman, Mohammad Shagrani, Eissa Faqeih, and Fowzan S Alkuraya. Bial-lelic mutations in tetratricopeptide repeat domain 26 (intraflagellar transport 56) cause severe biliary ciliopathy in humans. Hepatology, 71(6):2067–2079, jun 2020. doi: 10.1002/hep.30982.

14. Joanna Walczak-Sztulpa, Anna Wawrocka, Małgorzata Stańczyk, Karolina Pesz, Lech Du-darewicz, Sławomir Chrul, Ewelina Bukowska-Olech, Nina Wieczorek-Cichecka, Heleen H Arts, Machteld M Oud, Robert S’ migiel, Ryszard Grenda, Ewa Obersztyn, Krystyna H Chrzanowska, and Anna Latos-Bielenska. Interfamilial clinical variability in four polish families with cranioectodermal dysplasia and identical compound heterozygous variants in WDR35. American Journal of Medical Genetics. Part A, 185(4):1195–1203, apr 2021. doi: 10.1002/ajmg.a.62067.

15. Domenico Alvaro, Paolo Onori, Gianfranco Alpini, Antonio Franchitto, Douglas M Jefferson, Alessia Torrice, Vincenzo Cardinale, Fabrizio Stefanelli, Maria Grazia Mancino, Mario Strazzabosco, Mario Angelico, Adolfo Attili, and Eugenio Gaudio. Morphological and functional features of hepatic cyst epithelium in autosomal dominant polycystic kidney disease. The American Journal of Pathology, 172(2):321–332, feb 2008. doi: 10.2353/ajpath.2008.070293.

16. Tooba Quidwai, Jiaolong Wang, Emma A Hall, Narcis A Petriman, Weihua Leng, Petra Kiesel, Jonathan N Wells, Laura C Murphy, Margaret A Keighren, Joseph A Marsh, Esben Lorentzen, Gaia Pigino, and Pleasantine Mill. A WDR35-dependent coat protein complex transports ciliary membrane cargo vesicles to cilia. eLife, 10, nov 2021. ISSN 2050-084X. doi: 10.7554/{eLife}.69786.

17. Wei Wang, Tana S Pottorf, Henry H Wang, Ruochen Dong, Matthew A Kavanaugh, Joseph T Cornelius, Katie L Dennis, Udayan Apte, Michele T Pritchard, Madhulika Sharma, and Pamela V Tran. IFT-a deficiency in juvenile mice impairs biliary development and exacerbates ADPKD liver disease. The Journal of Pathology, 254(3):289–302, jul 2021. doi: 10.1002/path.5685.

18. Volker Bergen, Marius Lange, Stefan Peidli, F Alexander Wolf, and Fabian J Theis. Generalizing RNA velocity to transient cell states through dynamical modeling. Nature Biotechnology, 38(12):1408–1414, dec 2020. ISSN 1087-0156. doi: 10.1038/s41587-020-0591-3.

19. Gioele La Manno, Ruslan Soldatov, Amit Zeisel, Emelie Braun, Hannah Hochgerner, Viktor Petukhov, Katja Lidschreiber, Maria E Kastriti, Peter Lönnerberg, Alessandro Furlan, Jean Fan, Lars E Borm, Zehua Liu, David van Bruggen, Jimin Guo, Xiaoling He, Roger Barker, Erik Sundström, Gonçalo Castelo-Branco, Patrick Cramer, Igor Adameyko, Sten Linnars-son, and Peter V Kharchenko. RNA velocity of single cells. Nature, 560(7719):494–498, aug 2018. ISSN 0028-0836. doi: 10.1038/s41586-018-0414-6.

20. Jinbiao Chen, Ngan Ching Cheng, Jade A Boland, Ken Liu, James G Kench, D Neil Watkins, Sofia Ferreira-Gonzalez, Stuart J Forbes, and Geoffrey W McCaughan. Deletion of kif3a in CK19 positive cells leads to primary cilia loss, biliary cell proliferation and cystic liver lesions in TAA-treated mice. Biochimica et biophysica acta. Molecular basis of disease, 1868(4): 166335, apr 2022. doi: 10.1016/j.bbadis.2021.166335.

21. Luciane M Silva, Damon T Jacobs, Bailey A Allard, Timothy A Fields, Madhulika Sharma, Darren P Wallace, and Pamela V Tran. Inhibition of hedgehog signaling suppresses proliferation and microcyst formation of human autosomal dominant polycystic kidney disease cells. Scientific Reports, 8(1):4985, mar 2018. doi: 10.1038/s41598-018-23341-2.

22. Pamela V Tran, George C Talbott, Annick Turbe-Doan, Damon T Jacobs, Michael P Schon-feld, Luciane M Silva, Anindita Chatterjee, Mary Prysak, Bailey A Allard, and David R Beier. Downregulating hedgehog signaling reduces renal cystogenic potential of mouse models. Journal of the American Society of Nephrology, 25(10):2201–2212, oct 2014. doi: 10.1681/{ASN}.2013070735.

23. Yasushi Miura, Satoshi Matsui, Naoko Miyata, Kenichi Harada, Yamato Kikkawa, Masaki Ohmuraya, Kimi Araki, Shinya Tsurusaki, Hitoshi Okochi, Nobuhito Goda, Atsushi Miya-jima, and Minoru Tanaka. Differential expression of lutheran/BCAM regulates biliary tissue remodeling in ductular reaction during liver regeneration. eLife, 7, jul 2018. doi: 10.7554/{eLife}.36572.

24. Naoki Tanimizu, Yamato Kikkawa, Toshihiro Mitaka, and Atsushi Miyajima. 1-and *α*5-containing laminins regulate the development of bile ducts via *β1* integrin signals. The Journal of Biological Chemistry, 287(34):28586–28597, aug 2012. doi: 10.1074/jbc.M112.350488.

25. Frédéric Clotman, Patrick Jacquemin, Nicolas Plumb-Rudewiez, Christophe E Pierreux, Patrick Van der Smissen, Harry C Dietz, Pierre J Courtoy, Guy G Rousseau, and Frédéric P Lemaigre. Control of liver cell fate decision by a gradient of TGF beta signaling modulated by onecut transcription factors. Genes & Development, 19(16):1849–1854, aug 2005. doi: 10.1101/gad.340305.

26. Naoki Tanimizu, Atsushi Miyajima, and Keith E Mostov. Liver progenitor cells develop cholangiocyte-type epithelial polarity in three-dimensional culture. Molecular Biology of the Cell, 18(4):1472–1479, apr 2007. doi: 10.1091/mbc.E06-09-0848.

27. Aura D Urribarri, Patricia Munoz-Garrido, María J Perugorria, Oihane Erice, Maite Merino-Azpitarte, Ander Arbelaiz, Elisa Lozano, Elizabeth Hijona, Raúl Jiménez-Agüero, Maite G Fernandez-Barrena, Juan P Jimeno, Marco Marzioni, Jose J G Marin, Tatyana V Masyuk, Nicholas F LaRusso, Jesús Prieto, Luis Bujanda, and Jesús M Banales. Inhibition of metalloprotease hyperactivity in cystic cholangiocytes halts the development of polycystic liver diseases. Gut, 63(10):1658–1667, oct 2014. doi: 10.1136/gutjnl-2013-305281.

28. Adrien Guillot, Lucia Guerri, Dechun Feng, Seung-Jin Kim, Yeni Ait Ahmed, Janos Paloczi, Yong He, Kornel Schuebel, Shen Dai, Fengming Liu, Pal Pacher, Tatiana Kisseleva, Xuebin Qin, David Goldman, Frank Tacke, and Bin Gao. Bile acid-activated macrophages promote biliary epithelial cell proliferation through integrin *β6* upregulation following liver injury. The Journal of Clinical Investigation, 131(9), may 2021. doi: 10.1172/{JCI132305}.

29. Eleonora Patsenker, Yury Popov, Felix Stickel, Alfred Jonczyk, Simon L Goodman, and Detlef Schuppan. Inhibition of integrin alphavbeta6 on cholangiocytes blocks transforming growth factor-beta activation and retards biliary fibrosis progression. Gastroenterology, 135 (2):660–670, aug 2008. ISSN 1528-0012. doi: 10.1053/j.gastro.2008.04.009.

30. Sabrine Hassane, Wouter N Leonhard, Annemieke van der Wal, Lukas Jac Hawinkels, Irma S Lantinga-van Leeuwen, Peter ten Dijke, Martijn H Breuning, Emile de Heer, and Dorien Jm Peters. Elevated TGFbeta-smad signalling in experimental pkd1 models and human patients with polycystic kidney disease. The Journal of Pathology, 222(1):21–31, sep 2010. doi: 10.1002/path.2734.

31. B P Young, R A Craven, P J Reid, M Willer, and C J Stirling. Sec63p and kar2p are required for the translocation of SRP-dependent precursors into the yeast endoplasmic reticulum in vivo. The EMBO Journal, 20(1-2):262–271, jan 2001. doi: 10.1093/emboj/20.1.262.

32. Sorin V Fedeles, Xin Tian, Anna-Rachel Gallagher, Michihiro Mitobe, Saori Nishio, Se-ung Hun Lee, Yiqiang Cai, Lin Geng, Craig M Crews, and Stefan Somlo. A genetic interaction network of five genes for human polycystic kidney and liver diseases defines polycystin-1 as the central determinant of cyst formation. Nature Genetics, 43(7):639–647, jun 2011. ISSN 1546-1718. doi: 10.1038/ng.860.

33. Meritxell Huch, Craig Dorrell, Sylvia F Boj, Johan H van Es, Vivian S W Li, Marc van de Wetering, Toshiro Sato, Karien Hamer, Nobuo Sasaki, Milton J Finegold, Annelise Haft, Robert G Vries, Markus Grompe, and Hans Clevers. In vitro expansion of single lgr5+ liver stem cells induced by wnt-driven regeneration. Nature, 494(7436):247–250, feb 2013. doi: 10.1038/nature11826.

34. Darrell Andrews, Giorgio Oliviero, Letizia De Chiara, Ariane Watson, Emily Rochford, Kieran Wynne, Ciaran Kennedy, Shane Clerkin, Benjamin Doyle, Catherine Godson, Paul Connell, Colm O’Brien, Gerard Cagney, and John Crean. Unravelling the transcriptional responses of TGF-*β*: Smad3 and EZH2 constitute a regulatory switch that controls neuroretinal epithelial cell fate specification. The FASEB Journal, 33(5):6667–6681, may 2019. doi: 10.1096/fj.{201800566RR}.

35. Jenna Graham, Michael Raghunath, and Viola Vogel. Fibrillar fibronectin plays a key role as nucleator of collagen i polymerization during macromolecular crowding-enhanced matrix assembly. Biomaterials science, 7(11):4519–4535, nov 2019. doi: 10.1039/c9bm00868c.

36. Simon Cabello-Aguilar, Mélissa Alame, Fabien Kon-Sun-Tack, Caroline Fau, Matthieu Lacroix, and Jacques Colinge. SingleCellSignalR: inference of intercellular networks from single-cell transcriptomics. Nucleic Acids Research, 48(10):e55, jun 2020. doi: 10.1093/nar/gkaa183.

37. Mirjana Efremova, Miquel Vento-Tormo, Sarah A Teichmann, and Roser Vento-Tormo. CellPhoneDB: inferring cell-cell communication from combined expression of multi-subunit ligand-receptor complexes. Nature Protocols, 15(4):1484–1506, apr 2020. ISSN 1754-2189. doi: 10.1038/s41596-020-0292-x.

38. Jenny Z Kechagia, Johanna Ivaska, and Pere Roca-Cusachs. Integrins as biomechanical sensors of the microenvironment. Nature Reviews. Molecular Cell Biology, 20(8):457–473, aug 2019. ISSN 1471-0072. doi: 10.1038/s41580-019-0134-2.

39. Nasreen Akhtar and Charles H Streuli. An integrin-ILK-microtubule network orients cell polarity and lumen formation in glandular epithelium. Nature Cell Biology, 15(1):17–27, jan 2013. doi: 10.1038/ncb2646.

40. Carina Magdaleno, Trenton House, Jogendra S Pawar, Sophia Carvalho, Narendiran Ra-jasekaran, and Archana Varadaraj. Fibronectin assembly regulates lumen formation in breast acini. Journal of Cellular Biochemistry, 122(5):524–537, may 2021. doi: 10.1002/jcb.29885.

41. Hendrik A Messal, Silvanus Alt, Rute M M Ferreira, Christopher Gribben, Victoria Min-Yi Wang, Corina G Cotoi, Guillaume Salbreux, and Axel Behrens. Tissue curvature and api-cobasal mechanical tension imbalance instruct cancer morphogenesis. Nature, 566(7742): 126–130, feb 2019. ISSN 0028-0836. doi: 10.1038/s41586-019-0891-2.

42. Manoe J Janssen, Esmé Waanders, René H M Te Morsche, Ruoyu Xing, Henry B P M Dijkman, Jannes Woudenberg, and Joost P H Drenth. Secondary, somatic mutations might promote cyst formation in patients with autosomal dominant polycystic liver disease. Gastroenterology, 141(6):2056–2063.e2, dec 2011. ISSN 1528-0012. doi: 10.1053/j.gastro.2011.08.004.

43. Manoe J Janssen, Jody Salomon, René H M Te Morsche, and Joost P H Drenth. Loss of heterozygosity is present in SEC63 germline carriers with polycystic liver disease. Plos One, 7(11):e50324, nov 2012. doi: 10.1371/journal.pone.0050324.

44. Manoe J Janssen, Jody Salomon, Wybrich R Cnossen, Carsten Bergmann, Rolph Pfundt, and Joost P H Drenth. Somatic loss of polycystic disease genes contributes to the formation of isolated and polycystic liver cysts. Gut, 64(4):688–690, apr 2015. ISSN 0017-5749. doi: 10.1136/gutjnl-2014-308062.

45. Dustin J Flanagan, Nalle Pentinmikko, Kalle Luopajärvi, Nicky J Willis, Kathryn Gilroy, Alexander P Raven, Lynn Mcgarry, Johanna I Englund, Anna T Webb, Sandra Scharaw, Nadia Nasreddin, Michael C Hodder, Rachel A Ridgway, Emma Minnee, Nathalie Sphyris, Ella Gilchrist, Arafath K Najumudeen, Beatrice Romagnolo, Christine Perret, Ann C Williams, Hans Clevers, Pirjo Nummela, Marianne Lähde, Kari Alitalo, Ville Hietakangas, Ann Hedley, William Clark, Colin Nixon, Kristina Kirschner, E Yvonne Jones, Ari Ristimäki, Simon J Leedham, Paul V Fish, Jean-Paul Vincent, Pekka Katajisto, and Owen J Sansom. NO-TUM from apc-mutant cells biases clonal competition to initiate cancer. Nature, 594(7863): 430–435, jun 2021. ISSN 0028-0836. doi: 10.1038/s41586-021-03525-z.

46. D J Huels, L Bruens, M C Hodder, P Cammareri, A D Campbell, R A Ridgway, D M Gay, M Solar-Abboud, W J Faller, C Nixon, L B Zeiger, M E McLaughlin, E Morrissey, D J Winton, H J Snippert, J van Rheenen, and O J Sansom. Wnt ligands influence tumour initiation by controlling the number of intestinal stem cells. Nature Communications, 9(1):1132, mar 2018. doi: 10.1038/s41467-018-03426-2.

47. Arnout G Schepers, Hugo J Snippert, Daniel E Stange, Maaike van den Born, Johan H van Es, Marc van de Wetering, and Hans Clevers. Lineage tracing reveals lgr5+ stem cell activity in mouse intestinal adenomas. Science, 337(6095):730–735, aug 2012. doi: 10.1126/science.1224676.

48. Anne C Rios, Bianca D Capaldo, François Vaillant, Bhupinder Pal, Ravian van Ineveld, Caleb A Dawson, Yunshun Chen, Emma Nolan, Nai Yang Fu, 3DTCLSM Group, Felicity C Jackling, Sapna Devi, David Clouston, Lachlan Whitehead, Gordon K Smyth, Scott N Mueller, Geoffrey J Lindeman, and Jane E Visvader. Intraclonal plasticity in mammary tumors revealed through large-scale single-cell resolution 3D imaging. Cancer Cell, 35(4): 618–632.e6, apr 2019. ISSN 15356108. doi: 10.1016/j.ccell.2019.02.010.

49. Emilie Cornec-Le Gall, Ahsan Alam, and Ronald D Perrone. Autosomal dominant polycystic kidney disease. The Lancet, 393(10174):919–935, mar 2019. doi: 10.1016/S0140-6736(18)32782-X.

50. Alistair J Langlands, Axel A Almet, Paul L Appleton, Ian P Newton, James M Osborne, and Inke S Näthke. Paneth cell-rich regions separated by a cluster of lgr5+ cells initiate crypt fission in the intestinal stem cell niche. PLoS Biology, 14(6):e1002491, jun 2016. doi: 10.1371/journal.pbio.1002491.

51. H S Park, R A Goodlad, and N A Wright. Crypt fission in the small intestine and colon. a mechanism for the emergence of G6PD locus-mutated crypts after treatment with mutagens. The American Journal of Pathology, 147(5):1416–1427, nov 1995.

52. Naren P Tallapragada, Hailey M Cambra, Tomas Wald, Samantha Keough Jalbert, Diana M Abraham, Ophir D Klein, and Allon M Klein. Inflation-collapse dynamics drive patterning and morphogenesis in intestinal organoids. Cell Stem Cell, 28(9):1516–1532.e14, sep 2021. ISSN 19345909. doi: 10.1016/j.stem.2021.04.002.

## Bibliography

1. Pleasantine Mill, Paul J Lockhart, Elizabeth Fitzpatrick, Hayley S Mountford, Emma A Hall, Martin A M Reijns, Margaret Keighren, Melanie Bahlo, Catherine J Bromhead, Peter Budd, Salim Aftimos, Martin B Delatycki, Ravi Savarirayan, Ian J Jackson, and David J Amor. Human and mouse mutations in WDR35 cause short-rib polydactyly syndromes due to abnormal ciliogenesis. American Journal of Human Genetics, 88(4):508–515, apr 2011. doi: 10.1016/j.ajhg.2011.03.015.

2. Anna L Means, Yanwen Xu, Aizhen Zhao, Kevin C Ray, and Guoqiang Gu. A CK19(CreERT) knockin mouse line allows for conditional DNA recombination in epithelial cells in multiple endodermal organs. Genesis, 46(6):318–323, jun 2008. doi: 10.1002/dvg.20397.

3. Arnout G Schepers, Hugo J Snippert, Daniel E Stange, Maaike van den Born, Johan H van Es, Marc van de Wetering, and Hans Clevers. Lineage tracing reveals lgr5+ stem cell activity in mouse intestinal adenomas. Science, 337(6095):730–735, aug 2012. doi: 10.1126/science.1224676.

4. Jonathan Taylor and Robert J Tibshirani. Statistical learning and selective inference. Proceedings of the National Academy of Sciences of the United States of America, 112(25): 7629–7634, jun 2015. doi: 10.1073/pnas.1507583112.

5. Luigi Aloia, Mikel Alexander McKie, Grégoire Vernaz, Lucía Cordero-Espinoza, Niya Aleksieva, Jelle van den Ameele, Francesco Antonica, Berta Font-Cunill, Alexander Raven, Riccardo Aiese Cigliano, German Belenguer, Richard L Mort, Andrea H Brand, Magdalena Zernicka-Goetz, Stuart J Forbes, Eric A Miska, and Meritxell Huch. Epigenetic remodelling licences adult cholangiocytes for organoid formation and liver regeneration. Nature Cell Biology, 21(11):1321–1333, nov 2019. ISSN 1465-7392. doi: 10.1038/s41556-019-0402-6.

6. Gioele La Manno, Ruslan Soldatov, Amit Zeisel, Emelie Braun, Hannah Hochgerner, Viktor Petukhov, Katja Lidschreiber, Maria E Kastriti, Peter Lönnerberg, Alessandro Furlan, Jean Fan, Lars E Borm, Zehua Liu, David van Bruggen, Jimin Guo, Xiaoling He, Roger Barker, Erik Sundström, Gonçalo Castelo-Branco, Patrick Cramer, Igor Adameyko, Sten Linnarsson, and Peter V Kharchenko. RNA velocity of single cells. Nature, 560(7719):494–498, aug 2018. ISSN 0028-0836. doi: 10.1038/s41586-018-0414-6.

7. Volker Bergen, Marius Lange, Stefan Peidli, F Alexander Wolf, and Fabian J Theis. Generalizing RNA velocity to transient cell states through dynamical modeling. Nature Biotechnology, 38(12):1408–1414, dec 2020. ISSN 1087-0156. doi: 10.1038/s41587-020-0591-3.

8. Simon Cabello-Aguilar, Mélissa Alame, Fabien Kon-Sun-Tack, Caroline Fau, Matthieu Lacroix, and Jacques Colinge. SingleCellSignalR: inference of intercellular networks from single-cell transcriptomics. Nucleic Acids Research, 48(10):e55, jun 2020. doi: 10.1093/nar/gkaa183.

9. Helga Thorvaldsdóttir, James T Robinson, and Jill P Mesirov. Integrative genomics viewer (IGV): high-performance genomics data visualization and exploration. Briefings in Bioinformatics, 14(2):178–192, mar 2013. doi: 10.1093/bib/bbs017.

10. KD Salter, R M Roman, N R LaRusso, J G Fitz, and R B Doctor. Modified culture conditions enhance expression of differentiated phenotypic properties of normal rat cholangiocytes. Laboratory Investigation, 80(11):1775–1778, nov 2000. doi: 10.1038/labinvest.3780187.

11. Jesús M Banales, Tatyana V Masyuk, Pamela S Bogert, Bing Q Huang, Sergio A Gradilone, Seung-Ok Lee, Angela J Stroope, Anatoliy I Masyuk, Juan F Medina, and Nicholas F LaRusso. Hepatic cystogenesis is associated with abnormal expression and location of ion transporters and water channels in an animal model of autosomal recessive polycystic kidney disease. The American Journal of Pathology, 173(6):1637–1646, dec 2008. doi: 10.2353/ajpath.2008.080125.

12. Caleb A Dawson, Bhupinder Pal, François Vaillant, Luke C Gandolfo, Zhaoyuan Liu, Camille Bleriot, Florent Ginhoux, Gordon K Smyth, Geoffrey J Lindeman, Scott N Mueller, Anne C Rios, and Jane E Visvader. Tissue-resident ductal macrophages survey the mammary epithelium and facilitate tissue remodelling. Nature Cell Biology, 22(5):546–558, may 2020. doi: 10.1038/s41556-020-0505-0.

13. Nicholas T Younger, Mollie L Wilson, Anabel Martinez Lyons, Edward J Jarman, Alison M Meynert, Graeme R Grimes, Konstantinos Gournopanos, Scott H Waddell, Peter A Tennant, David H Wilson, Rachel V Guest, Stephen J Wigmore, Juan Carlos Acosta, Timothy J Kendall, Martin S Taylor, Duncan Sproul, Pleasantine Mill, and Luke Boulter. In vivo modeling of patient genetic heterogeneity identifies new ways to target cholangiocarcinoma. Cancer Research, jan 2022. ISSN 0008-5472. doi: 10.1158/0008-5472.{CAN}-21-2556.

