## Supplemental Materials, Methods and Acknowledgements for "Primary cilia loss promotes reactivation of morphogenesis and cyst-fission through a deregulated TGF*β*-ECM-Integrin axis in polycystic liver disease"

[1]MRC Human Genetics Unit, Institute of Genetics and Cancer, University of Edinburgh, Edinburgh, UK, EH4 2XU
[2]School of Informatics, University of Edinburgh, Edinburgh, UK, EH8 9AB
[3]Department of Liver and Gastrointestinal Diseases, Biodonostia Health Research Institute, University of the Basque Country, San Sebastian, Spain, 20014
[4]Department of Gastroenterology and Hepatology, Radboud University, Nijmegen Medical Center, 6525 GA Nijmegen, The Netherlands
[5]Centre for Inflammation Research, Queens Medical Research Institute, University of Edinburgh, Edinburgh, UK EH16 4TJ
[6]National Institute for the Study of Liver and Gastrointestinal Diseases (CIBERehd, "Instituto de Salud Carlos III"), Spain
[7]Department of Biochemistry and Genetics, School of Sciences, University of Navarra, Pamplona, Spain, 31008
[8]IKERBASQUE, Basque Foundation for Science, Bilbao, Spain, 48009
[*]These authors contributed equally to this work

**polycystic disease | cilia | cholangiocyte | morphogenesis | extracellular matrix | Integrin | TGF$\beta$**

### Online Materials and Methods

#### Human Tissue:

Human normal liver tissue and liver from patients with liver cysts were obtained from the NRS BioResource and Dr Tim Kendall, NHS consultant pathologist, confirmed pathology. JPH Drenth, Department of Gastroenterology and Hepatology, Institute kindly provided samples from patients with PCLD with known genetics for Molecular Life Sciences, Radboud University Medical Center. All tissues were collected under informed consent or were archival diagnostic specimens. Details of human tissue is summarised in **Supplementary table 1**.
